# Beyond the skin barrier: commensal *S. epidermidis* imprint systemic immunity to invasive biofilm infection

**DOI:** 10.64898/2026.01.08.698386

**Authors:** Pia Fehrenbach, Elian M. A. Kuhn, Lena Gens, James Tapia-Dean, Javier Rangel-Moreno, Alisa Hangartner, Puk Kwant, Stephan Zeiter, Esther C. de Jong, Gowrishankar Muthukrishnan, T. Fintan Moriarty

## Abstract

*Staphylococcus epidermidis*, a dominant human skin commensal from early life, can transition to an opportunistic pathogen, including invasive, biofilm-associated infections linked to medical devices. Neonatal exposure to skin commensals induces a lifelong immunological imprint in the skin, characterized by immunoregulatory responses. We therefore hypothesized that early life exposure to *S. epidermidis* influences immune responses to invasive biofilm-associated infections later in life.

Using a mouse model of biofilm-related *S. epidermidis* bone infection, we show that neonatal and adult skin colonization altered the immune response to the subsequent infection in adulthood. Neonatal colonization led to increased NK cells and neutrophils compared to no colonization, along with reduced Tregs and Th1 cells, and consistent increase in immune checkpoint receptor PD-1^+^ Tregs, T effector, Th1 and Th2 cells across infected bone marrow, blood and spleen. These PD-1-related immune modulations were absent in the adult-colonized group, which had the highest numbers of Tregs, Th1 and Th2 cells of all groups.

These findings reveal that early exposure to commensal bacteria strongly impacts the response to invasive infection later in life. Notably, the response depends on the timing of previous exposure. Neonatal colonization drives T cell modulation, resembling neonatal immunity, while adult-colonization increases specific T cell abundance. These differences highlight the essential role of skin colonization in shaping the quality of pathogen immunity to protect against invasive, biofilm-associated infection later in life, emphasizing that immunological studies using uncolonized animal models may not fully capture human immune dynamics.

## Introduction

*Staphylococcus epidermidis* is a Gram-positive commensal bacterium that colonizes human skin and mucous membranes from birth ^1,2^. Similar to other commensal bacteria of the skin microbiome, *S. epidermidis* plays a critical role in maintaining skin barrier homeostasis, microbial balance, and protection against pathogens through direct skin interactions, colonization resistance, and modulation of immunity ^3-5^.

Immune responses to skin commensals are generally biased toward immunoregulation and tolerance in the neonate, as shown in mice ^2,6-8^. These neonate-exposure-driven responses show local expansion and recruitment of regulatory T cells (Treg) to the skin ^9,10^, increased IL-1 signaling, and suppression of Th17 responses ^11^. When *S. epidermidis* colonization is experimentally delayed to adulthood, the neonatal Treg-dominated response disappears, rather, a CD8^+^ T cell and effector T cell-dominated response occurs. Thus, inducing skin inflammation rather than tolerance ^10-12^.

Whilst colonization-induced immunoregulatory responses seem relatively well characterized for skin commensals, it remains largely unexplored how prior colonization by commensal bacteria impacts opportunistic, invasive, biofilm-associated infections caused by these bacteria later in life. *S. epidermidis* has emerged as an opportunistic pathogen in patients with indwelling medical devices (e.g., catheters, pacemakers, and ventilation tubes), whereby it accounts for up to 50 % of cases of late-developing infections ^13,14^, which may be particularly relevant against a background of an increasingly aging population ^15^. Its ability to form biofilms on medical devices significantly reduces bacterial clearance by antibiotics and the immune system ^16-19^. Further complicating treatment is the emergence of antibiotic resistance ^20,21^. Whether early-life immune imprinting, previously described in the skin also influences immune responses during invasive *S. epidermidis* infections that occur later in life remains to be determined.

This study aimed to test the hypothesis that early-life skin colonization with *S. epidermidis* imprints a lasting systemic immune response, thereby altering immunity to invasive bone infection later in life by shaping the balance between infection control and immune tolerance. This would also provide insight into laboratory animal studies of host-pathogen interactions where animal colonization may need to be more controlled.

## Results

### Immune Profiling Reveals Long-Term Imprint of Colonization on Subsequent Infection

The effect of neonatal skin exposure on subsequent *S. epidermidis* deep bone infection was determined under controlled conditions in a well-defined mouse tibial pin model using flow cytometry. To compare the overall phenotypical similarities of immune cells between colonization groups, clusters based on marker intensity and combination were visualized as UMAPs. In the bone marrow (infection site) it showed that neonatally colonized mice (*n*=8) displayed a visually distinct immune landscape compared to both non-colonized (*n*=8) and adult-colonized mice (*n*=6) (**Figure 1A**). Variations in group size resulted from the exclusion of animals. Unsupervised clustering with PhenoGraph, combined with differential abundance analysis using edgeR, identified significant differences between neonatal-colonized and non-colonized mice (29 clusters) and adult-colonized mice (10 clusters) (**Table S1**). The response of both neonatal and adult colonized mice, based on cluster analysis, was primarily characterized by an increase in NK cells, compared with non-colonized mice (5 NK-related clusters increased). Adult-colonized mice had increased T and B cells, compared to neonatal-colonized mice (**Figure 1B**). Equivalent data for spleen and blood are shown in the supplementary **Figures S1** and **S2**.

**Figure 1:**
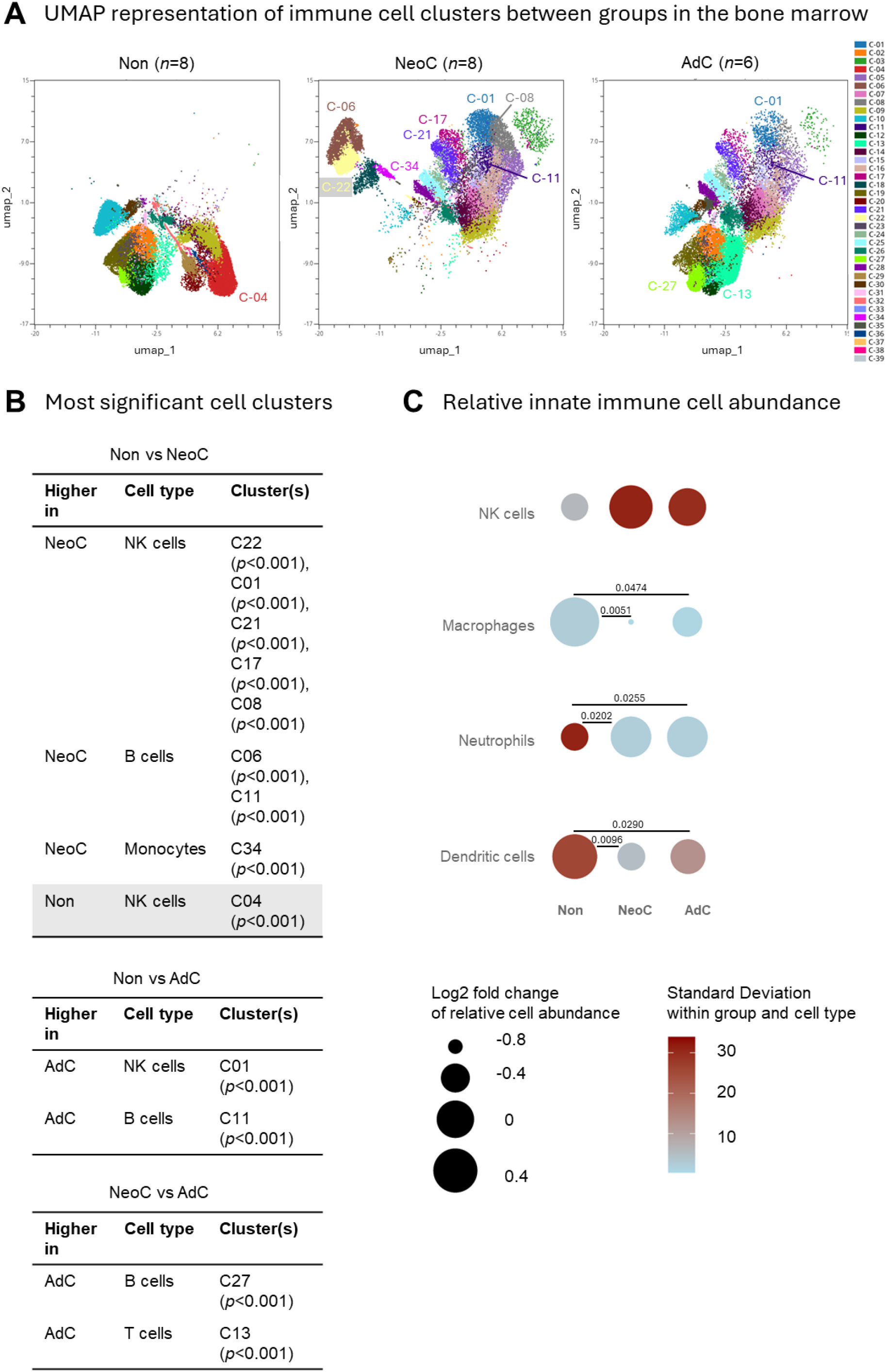
Comparison of immune cell clusters and innate immune cell abundance in bone marrow of colonized and non-colonized mice. **A)** UMAP representative of all significantly different clusters from non-colonized (Non, *n* = 8), neonatal-colonized (NeoC, *n* = 8), and adult-colonized mice (AdC, *n* = 6). **B)** The highest significantly different clusters above log(p-value) of 3.5 from each comparison obtained with Welch’s t-test are listed in the tables comparing the groups. **C)** The difference between innate cell percentage in Non, NeoC, and AdC groups is compared for natural killer cells (NK), macrophages, neutrophils, and dendritic cells (DC) in the bone marrow. It is visualized by plotting the log_2_ fold change of each group mean relative to the overall mean of the three group means (circle area size), with color indicating the within-group standard deviation. Flow cytometry data from individual mice were calculated (Non: n = 8; NeoC: n = 8; AdC: n = 6). Statistical analyses were performed with one-way ANOVA with Tukey’s multiple comparisons test or non-parametric with Kruskal-Wallis test - only significant p-values shown.

For marker specific cell population investigation, conventional flow cytometry gating was performed. Cell populations, presented as log_2_ fold change of each groups mean compared to the mean between groups (**Figure 1C**), once again revealed that NK cells were elevated in colonized mice (non-significant), mainly the neonatal-colonized group, but also in the adult-colonized group. Upon examination at the individual mouse level (**Figure S3**), a bimodal distribution was observed, with a subset of mice displaying particularly high NK cell enrichment. Both neonate (p = 0.0202) and adult (p = 0.0255) colonization were also associated with a marked and consistent increase in neutrophils, but a decrease in macrophages (neonatal p = 0.0051; adult p = 0.0474) and dendritic cells (neonatal p = 0.0096; adult p = 0.0290) compared with non-colonized mice (Figure 1C).

The differences seen between cluster and gating results arise because cluster analysis incorporates marker intensities to define subpopulations within each cell type, whereas conventional gating groups all marker-positive cells together regardless of intensity. Collectively, these results show distinct alterations in immune cell composition comparing colonized with non-colonized groups, with the most prominent changes observed in NK cells, neutrophils, macrophages, and dendritic cells.

### T Cell Landscape Reveals Long-Term Imprint of Colonization Timing in Bone Marrow

Since the cluster analysis revealed significant differences in T cell populations between neonatal- and adult-colonized groups, we next examined T cell subsets in greater detail to assess the impact of colonization timing. The T cell cluster-based UMAP from the bone marrow (**Figure 2A**), revealed substantial differences. The neonatal- and adult-colonized groups displayed diminished amounts of cells in clusters C04 and C11, compared with the non-colonized group (Figure 2A). Both are clusters of CD3^+^CD4^+^IL4^+^ Th2 cells (**Figure 2B**). Neonatal-colonized mice had a decrease in Th2 cell clusters C14, C15, and C17, compared with the adult-colonized mice. Tregs in cluster C18 were also diminished in the neonatal-colonized group, but only in comparison with the adult group. Equivalent data for spleen and blood are shown in **Figure S4** and **S5**. Cells from Th2 cell clusters were increased in the spleen of both colonized groups, while Treg clusters were decreased in colonized groups. The identities of all significant clusters are shown in supplementary **Table S2**.

**Figure 2:**
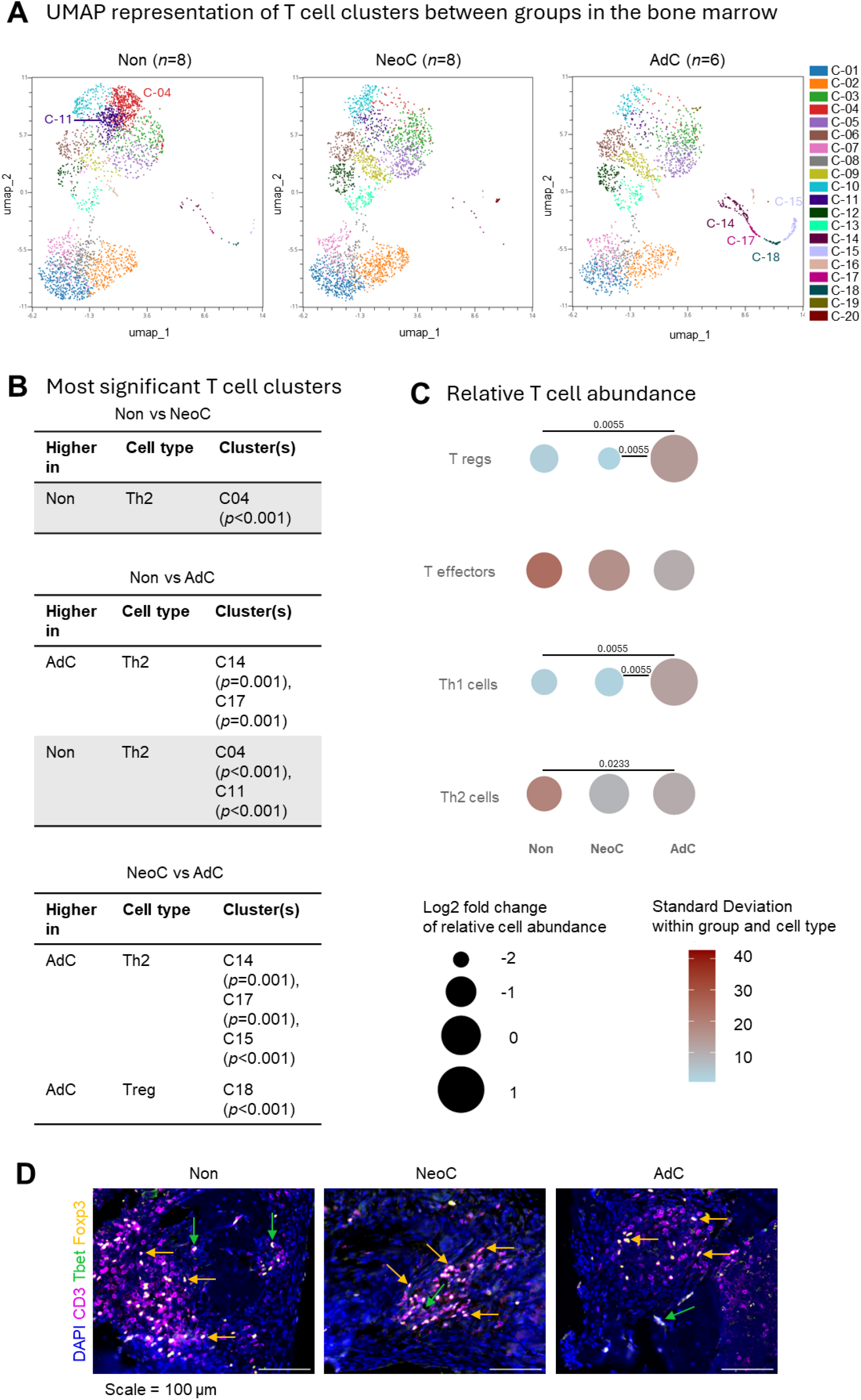
Comparison of T cell clusters in bone marrow of colonized and non-colonized mice. **A)** UMAP representative of all significantly different T cell clusters from non-colonized (Non: n = 8), neonatal-colonized (NeoC: n = 8), and adult-colonized mice (AdC: n = 6). **B)** The highest significantly different clusters (Log p-value above 2.5) obtained using Welch’s t-test are listed in the table for Non and NeoC, Non and AdC and NeoC and AdC comparison. **C)** The difference between T cell percentage in Non, NeoC, and AdC groups is compared for bone marrow’s natural killer cells (NK), macrophages, neutrophils, and dendritic cells (DC). It is visualized by plotting the log_2_ fold change of each group mean relative to the overall mean of the three group means (circle area size), with color indicating the within-group standard deviation. Flow cytometry data from individual mice (Non, n = 8; NeoC, n = 8; AdC, n = 6). Statistical analyses performed using one-way ANOVA with Tukey’s multiple comparisons test or non-parametric with Kruskal-Wallis test - only significant p-values shown. **D)** Immunofluorescence was performed on decalcified infected tibia with surrounding soft tissue of non-(Non: n= 1), neonatal-(NeoC: n= 1) and adult-colonized mice (AdC: n= 1). Representative 100X-magnification pictures show the spatial location of CD3^+^ T cells, Th1 cells CD3^+^Tbet^+^ and TregsCD3^+^ Foxp3^+^ detected with specific antibodies, close to the pinhole. Green arrows point to CD3^+^Tbet^+^ Th1 cells, while yellow arrows point to CD3^+^ Foxp3^+^ Tregs cells. Scale = 100 µm

In contrast to the cluster analysis, the focused flow cytometry data from the bone marrow revealed that the neonatal-(non-significant) and adult-colonized (p = 0.0233) mice displayed elevated Th2 cells compared with non-colonized mice (**Figure 2C** and **Figure S6** for mouse-specific data). The most significant impact of adult colonization was a substantial increase in Tregs (p = 0.0055) and Th1 cells (p = 0.0055), particularly compared with the neonatal-colonized group. Comparisons between Th17 (CD3^+^, Il-17a^+^) and CD8^+^ T cells (CD3^+^, CD8^+^) showed no differences between the groups, but B cells (CD3^-^, CD19^+^) were higher in the bone marrow of non-colonized than colonized mice (**Figure S7**). Immunofluorescence of lesions inside the infected bone marrow or surrounding soft tissue showed abundant CD3^+^Foxp3^+^ Tregs and scarce CD3^+^Tbet^+^ Th1 cells after challenge with *S. epidermidis* in all the experimental mice (**Figure 2D**).

Overall, the analysis revealed that the timing of colonization corresponded with discrete shifts in T cell subset distribution, with increased Tregs, Th1 and Th2 cells for adult-colonized mice.

### PD1 Expression on T Cells Reveals Long-Term Imprint of Colonization Timing in bone marrow

Since functional activation, rather than abundance alone, defines T cell responses, Programmed cell Death protein 1 (PD-1) expression was analysed to assess activation and exhaustion patterns among T cell subsets. It showed a broad increase in PD-1^+^ cells in neonatal-colonized mice (**Figure 3A**, and mouse specific data in **Figure S8**). Specifically, neonatal-colonized mice had increased Tregs, effector T cells and Th1 cells with PD-1 expression compared with non-colonized (*p = 0*.*0092*, effector T cells) and adult colonized mice (non-significant). PD-1 expression in blood and spleen is shown in supplementary Figure S8, with PD1^+^ Th1 cells and Tregs again increased in the neonatal-colonized group.

**Figure 3:**
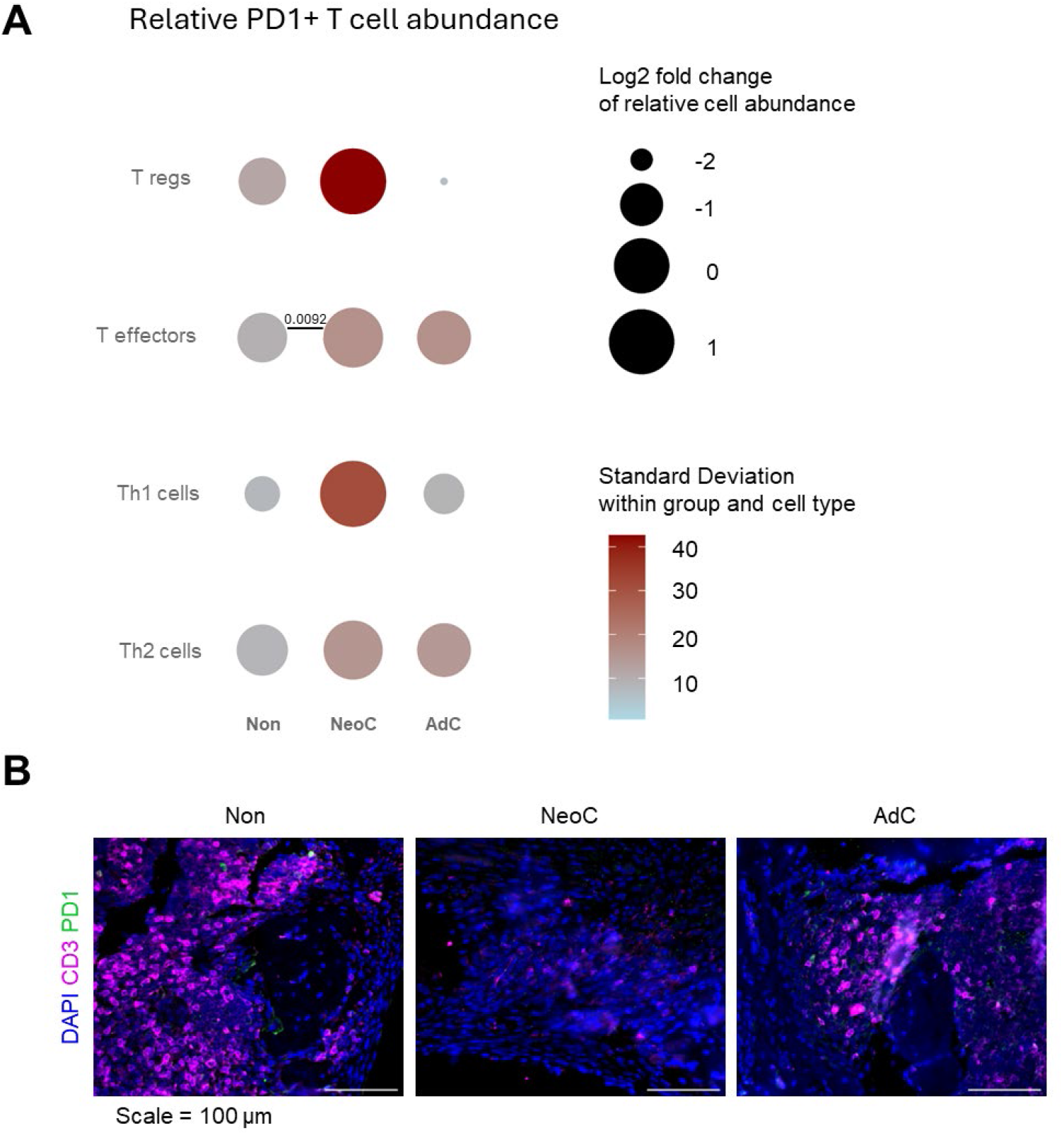
T cell subsets and PD-1 expression within T cell subsets in bone marrow. **A**) Comparative analysis of the difference between cell percentage expressing the exhaustion marker PD-1 in non-colonized (Non), neonatal colonized (NeoC) and adult-colonized (AdC) groups for bone marrow’s T regulatory cells (Tregs), T effectors, T helper 1 (Th1) and T helper 2 cells (Th2). It is visualized by plotting the log_2_ fold change of each group mean relative to the overall mean of the three group means (circle area size), with color indicating the within-group standard deviation. Calculation data from individual mice (Non, *n* = 8; NeoC, *n* = 8; AdC, *n* = 6). Statistical analyses performed using one-way ANOVA with Tukey’s multiple comparisons test or non-parametric with Kruskal-Wallis test - only significant p-values shown. **B)** Immunofluorescent staining was performed on decalcified infected tibia and surrounding soft tissue of non-(Non: n= 1), neonatal-(NeoC: n= 1) and adult-colonized mice (AdC: n= 1). Representative 100X-magnification pictures show the spatial location of CD3^+^PD-1^-^ T cells, and PD1^+^ cells near the pinhole. Scale = 100 µm

We next determined whether PD-1 expression levels reflected an active (low expression) or an exhausted T cell phenotype (high expression). Cells expressed low, intermediate and high levels of PD-1 (**Figure S9**). All tested PD1^+^ T cells from neonatal-colonized mice had predominantly low (L) PD-1 expression, some intermediate (I) PD-1 expression and almost no high (H) PD-1 expression. Detailed, Th2 (89% L; 10% I; 1% H) had lowest expression, followed by T effector cells (83% L; 16% I; 1% H), Tregs (57% L; 15% I; 28% H) and Th1 cells (44% L; 32% I; 24% H).

Consistent with the low percentage of T cells expressing high levels of PD-1, immunofluorescent staining of lesions at the infection site with CD3 and PD1 antibodies showed the CD3^+^PD1^+^ T cells were present but low in all the experimental groups.

These data show that neonatal colonization is associated with an overall increase in PD-1^+^ T cells characterized predominantly by low PD-1 expression levels across subsets.

### Prior Skin Colonization Has Limited Impact on Osteolysis and Infection Outcomes

To complement the immunological analyses, infection clearance and osteolysis were evaluated using histological, radiographic and bacteriological analysis. Immunofluorescence with a Staphylococcal antibody, confirmed the presence of bacteria in all mice (**Figure 4A**). In B&B-stained slides, neonatal-colonized and non-colonized mice displayed a significant accumulation of granulocytes along pin implantation site (Figure 4A, asterisks). Lymphocyte infiltration was more modest in the mouse colonized at neonatal age compared to both adult- and non-colonized mice (empty squares).

**Figure 4:**
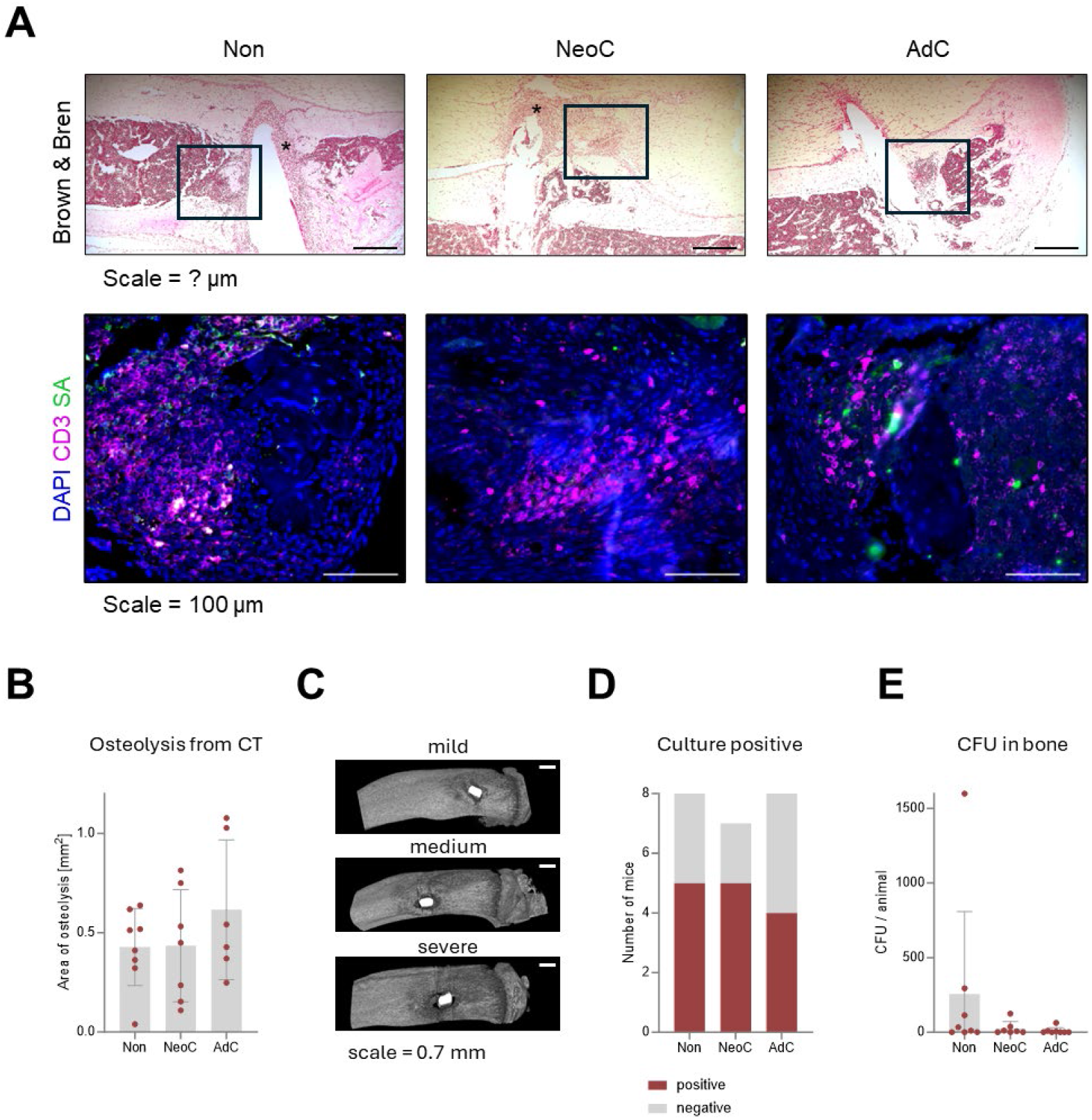
*S. epidermidis* culture positivity, bacterial counts, and osteolysis after euthanasia. **A)** Brown & Bren modified Gram staining and immunofluorescence in decalcified infected tibia with surrounding soft tissue of non-(Non: n= 1), neonatal-(NeoC: n= 1) and adult-colonized mice (AdC: n= 1). Black empty squares in representative 25x magnification pictures show areas with abundant immune cell infiltration, while asterisks depict infiltration by neutrophils in areas with pins coated with *S. epidermidis*. Scale = 100 μm. Representative 100X-magnification pictures show the spatial location of CD3^+^ T cells and *S. epidermidis*, detected with a *S. aureus* antibody, in the infected bones and surrounding soft tissue. **B)** MicroCT at euthanasia was used to quantify osteolysis areas in each animal. Data shown are from individual mice (Non, n = 8; NeoC, n = 7; AdC, n = 6). **C)** Representative microCT images visualizing osteolysis seen within each group in B. Scale = 0.7mm. **D)** Number of culture-positive mice as proportion of all mice within each group for all tissues combined. **E)** Colony forming units (CFU) were quantified in the bone at euthanasia for each animal per colonization group. Data shown are from individual mice (Non, n = 8; NeoC, n = 7; AdC, n = 8). Statistical was calculated with one-way ANOVA with Tukey’s multiple comparisons test.

Osteolysis surrounding the implants was evaluated at euthanasia (**Figure 4B**). Although osteolysis was higher on average in the adult-colonized group, the difference was not statistically significant, mainly due to the high variability observed between animals in each group (**Figure 4C**). There was no trend in osteolysis perimeter, nor was there a clear correlation between bacterial load (CFU) and osteolysis (**Figure S10A** and **B**).

The culture positivity rate is shown in **Figure 4D**, revealing that approximately half of the animals remained culture positive at euthanasia across all groups (neonatal-colonized: 5/7; non-colonized: 5/8, adult-colonized: 4/8). Culture positive mice showed low CFU counts and no statistically significant differences between the groups, though there was a trend with increased CFUs for some non-colonized mice in the bone (**Figure 4E, Figure S10C**), despite having the same inoculum (**Figure S10D**).

In summary, despite marked immune differences between groups, infection clearance and bone pathology were overall similar at the timepoint of euthanasia, with variability in bacterial load and osteolysis among individual animals. These bacteriological analyses were included to provide an overview of infection status at euthanasia, to better understand the immunological parameters. Despite the relatively limited differences in infection outcome, the marked differences in immune responses represent an important advance in our understanding of colonization effect on the immune response to infection.

## Discussion and conclusion

This study investigated whether skin colonization with *S. epidermidis* influences immune responses to subsequent invasive, biofilm-associated infection and, further, whether colonization during the neonatal period elicits distinct immune outcomes compared to colonization in adulthood.

One of the main findings that was impacted by prior colonisation (neonate or adult) was the response of the innate immune cell populations. Several NK cell clusters were elevated in previously colonized animals (both adult and neonatal). NK cells are innate lymphocytes with direct cytotoxic activity against infected or transformed cells ^22^ and have been shown to be present in osteomyelitic lesions in mice ^23^ and human patients ^24^. Previous studies have shown that NK cells do not contribute to a reduction in bacterial load in septic arthritis mouse model, but play a role in minimising the pathological appearance of arthritis, suggesting they have an important role in limiting inflammation-driven tissue damage ^24^. Further functional assessment, including cytokine secretion profiling, will be required to define the precise mechanism underlying this expansion.

Neutrophil infiltration is a well-recognized histopathological hallmark used in the diagnosis of acute osteomyelitis ^25^. Consistent with this, colonized mice exhibited elevated neutrophil numbers in the current study, reflecting the innate response to bacterial bone infection. The delayed or reduced neutrophil recruitment observed in non-colonized mice is unlikely to be substantially influenced by memory cell compartments, as neutrophils can independently sense and respond to pathogen-derived signals. Given that neutrophils are required for NK cell maturation and function in mice ^26^ the correlation between reduced NK cell numbers and diminished neutrophil recruitment may be mechanistically linked. However, the precise reason for the delay in non-colonized mice remains to be determined. In contrast to the increases in NK cells and neutrophils, macrophages and DCs were reduced in previously colonized mice. In a previous study with *S. epidermidis* bone infection without prior colonization, early innate responses were characterized by a rapid macrophage influx that resolved within two weeks ^27^. In the present study, prior colonization led to a reduction in macrophage counts compared with non-colonized mice three weeks post-inoculation, potentially reflecting a more rapid resolution of the inflammatory phase or a diminished induction of macrophage recruitment. Finally, bone marrow DCs in colonized mice were decreased. DCs are a bridge between innate and adaptive immunity and develop in the bone marrow before migrating to secondary lymphoid organs such as the spleen for antigen presentation to T cells ^28^. Thus, it is possible that the induction of adaptive immunity is enhanced at the time of euthanasia in previously colonized mice. Enrichment of bone marrow DCs in colonized mice suggests that there will be an increased pool of antigen-presenting cells that contribute to the induction of optimal T cells responses to clear *S. epidermidis* in the infected bones ^28^. However, additional studies will be necessary to determine if the change in number has an impact on functionality of these cells.

The timing of skin colonisation in neonates has previously been described and shown to involve local expansion and recruitment of Tregs to the skin accompanied by IL-1 and Th17 responses ^2,5,9-12^. In general, Treg development is more likely to occur during neonatal immune responses than during adult immune responses ^29^, and Treg generation happens mostly in peripheral tissue upon reaction to commensals ^30^. The findings of our study advance our understanding of this phenomenon, by showing that, in more invasive bone infection, adult-colonized mice displayed the more drastic increase in Tregs in bone marrow compared to the other groups. Given that T cell progenitors freshly derived in the bone marrow do not express CD4, CD8 or CD3 receptors, the T cells (including CD3^+^Foxp3^+^ Tregs) detected in our infected mice must be mature Tregs recruited to the site of infection ^31^. These high Treg numbers in adult-but not in neonatal-colonized mice contrast with the skin homeostasis studies, ^2,6,12^. Humans (colonized naturally with *S. epidermidis* neonatally) also show increased Tregs in osteomyelitis ^32^. The influx of Tregs in adult mice in invasive bone infection suggests that, in our model, the response to infection towards an immunoregulatory response might be more related to recency of exposure (2 weeks in the adult groups vs 9 weeks in neonates) rather than an immune imprinting in neonates, at least in relation to Tregs. However, to conclusively exclude this possibility, further investigations are needed, and the timing of the surgery should be postponed to a later stage. Alternatively, the Treg response might be influenced by distinct niches within the body, in which skin and subcutaneous tissue deliver signals different to the ones found in the bone marrow. Higher Treg abundance in adult-colonized mice could also suggest less effective bacterial clearance compared to neonatal-colonized mice.

The increase in Th1 and Th2 cells in adult-colonized mice reveals a balanced immune response. Th1 cells contribute to intracellular bacterial clearance, while the pronounced Th2 response may promote tissue repair and modulate Th1-driven tissue inflammation to prevent tissue damage. The neonatal-colonized immune response, marked by scarce Th1 cells and abundant Th2 cells, mirrors typical neonatal immune characteristics prioritizing Th2 cells to avoid inflammation from early microbial exposure ^33^. However, not only T cell numbers play a role for a balanced immune response, but also modulation of T cell activation. One of the more consistent findings, and also the main factor affected by neonatal exposure and absent in adult exposed or non-exposed animals, was in relation to PD-1. PD-1 is an immune checkpoint receptor expressed on the surface of T cells, B cells, and some myeloid cells ^34,35^. It downregulates immune responses to maintain self-tolerance and prevent autoimmunity. Importantly, early PD-1 expression suggests cell stimulation, while high levels of PD-1 at late stages of a chronic immune reaction likely correlate with impaired effector T cell functions ^36^. We found elevated numbers of all tested T cell subtypes, including Tregs, T effector, Th1, and Th2 cells expressing PD-1, suggesting enhanced functional modulation typical of neonatal immune adaptation ^33^. As PD-1 expression was mostly low or intermediate for PD-1^+^ T cells in neonatal-colonized mice, PD-1 is likely more indicative of high cell activity than cell exhaustion, which was confirmed by the lack of detection by immunofluorescence of PD-1^+^CD3^+^ T cells at the infection site. Therefore, despite low Treg and Th2 cell counts, PD-1^+^ T cell activation reveals a balanced response that incorporates effector and regulatory activity persisting into adulthood. Activated CD44^+^ Tregs were also observed in the skin of neonatal-colonized mice, as previously reported ^2^, and our findings confirm Tregs can migrate into infected bones.

Bacterial quantification and histology showed that in all groups, the infection was largely under control in all animals, with little to no CFU detected and immunofluorescence showing low bacterial signal. This model, suitable for bone infection study ^37^, has limitations because a strong immune clearance of *S. epidermidis* might not fully mirror human infection dynamics ^38,39^. Given the evident differences in immune responses observed among groups in this study, the relatively mild variations in infection outcomes between groups do not dictate these responses; instead, they reflect a true timing effect of exposure. The impact of skin colonization status should also be considered in immune-related animal experiments as humans become colonized directly at birth and animal studies are mostly done under sterile conditions. Lately, housing and pre-operative conditions for rodent animal studies have become more regulated and cleaner, which is beneficial for reproducibility and animal welfare ^40^. However, specific murine housing conditions can influence the microbiome and subsequent study outcomes ^41^.

Limitations of this study include the absence of longitudinal kinetics assessments for colonization dynamics between groups and control groups without infection; however, these were beyond the primary focus on the effects of colonization on bone infection. Moreover, at this point we cannot conclusively demonstrate that the immune phenotypes observed in adult-colonized animals resulted just from colonization-driven imprinting rather than being also influenced by transient immune priming, as colonization and infection were within 2 weeks. Further, although colonization was verified per cage with skin swabs and faeces samples at time of surgery, it was not confirmed for every individual animal, to reduce disturbance for the animals. To sum up, molecular approaches to confirm colonization, additional timepoints of euthanasia and cytokine patterns should be included in future studies.

In conclusion, this study demonstrates that prior microbial exposure via the skin critically shapes systemic immune responses to invasive infection. Using a mouse model, we show that non-colonized mice exhibit reduced recruitment of innate and adaptive immune cells and delayed immune regulation at infection sites. In contrast, neonatal colonization induces a balanced, functionally active immune profile characterized by regulated innate responses and heightened T cell activation, resembling early-life immunity. Adult colonization, however, promotes enhanced immune regulation through expansion of T cell populations, particularly Tregs, rather than activation. These findings reveal that the age at which colonization occurs “locks in” distinct immune states, with long-term consequences for systemic immunity.

Importantly, this work highlights the systemic impact of skin colonization beyond local tissue health and underscores the role of early microbial exposure in programming immune balance and T cell function. Such insights have broad implications for animal studies investigating vaccination strategies and other immunological interventions. Moreover, understanding the impact of PD-1, the PD-1 pathway and related immune checkpoints may open avenues for directed therapies or immunomodulatory strategies that prevent invasive, biofilm-associated infections without reliance on new antibiotics.

## Materials and methods

### Study design

48 C57BL/6J mice (24 male, 24 female) were divided into non-colonized, neonatal-colonized or adult-colonized group (Figure S11). Neonatal colonization with *S. epidermidis* was performed at 1 week of age and adult colonization at 8 weeks after birth. Subsequently, at 10 weeks of age, a pin inoculated with *S. epidermidis* was implanted in the right proximal tibia. Animals were euthanized 3 weeks later. Eight non-colonized (3 m/5 f), seven neonatal-colonized (3 m/4 f) and eight adult-colonized mice (4 m/4 f) were used for bacteriological outcomes and MicroCT image processing. Eight non-colonized (4 m/4 f), eight neonatal-colonized (4 m/4 f) and six adult-colonized mice (3 m/3 f) were reserved for flow cytometry analysis. One male per group was processed for histology.

### Study design

This study was approved by the local ethical IACUC and performed under Swiss animal protection law, approved license numbers: GR/14/2023, GR/31E/2023.

### Animals and housing

C57BL/6J breeding pairs (Charles River, Sulzfeld, Germany) and their litters were housed in IVC cages (BSL2 for colonized animals) with aspen bedding, a plastic house, nesting material and wooden gnawing material. They received extruded food (3436 Haltung Extrudat, Kliba Nafag, Kaiseraugst, Switzerland) and autoclaved water ad libitum. Mice were weaned at 3 weeks after birth, then separated by sex into small groups (2–4 per cage), with single housing only in exceptions. Parents were considered colonized after neonatal colonization of their litter, and their subsequent litter was solely used for the neonatal-colonized group. Post-weaning identification was done by toe-tattooing.

### Bacterial Preparation

*S. epidermidis* strain 103.1 (CCOS 1152), isolated from a bone infection, was genetically modified to express mCherry2 and chloramphenicol resistance (plasmid pJL74-2wmcherry-CAM, provided by Tiffany Scharschmidt group) by electroporation as previously described ^42^, using Tryptic Soy Broth (TSB, Oxiod, Basel, Switzerland). Plasmid insertion was confirmed by fluorescence microscopy. The antigen 2w was included for antigen-specific immune detection ^12^, but data are not shown due to low detection.

### Skin colonization

Skin colonization was performed according to established protocols ^2,12^. Briefly, planktonic *S. epidermidis*, was cultured in 20 mL TSB (10 µg/mL chloramphenicol) for 40 h at 37° C and 100 rpm, washed twice with 10 mL Phosphate Buffered Saline (PBS, Sigma, Basel, Switzerland) by centrifugation (1320 rcf, 7 min), sonicated 3 min at 40 kHz, and OD_600_ adjusted to 0.5. The bacterial suspension was applied to the back skin using sterile cotton swabs for 10 s under light pressure. Fur was shaved in 3-week-old mice before topical application. Colonization was repeated three times (2-day intervals).

### Preparation of inoculated pins

The murine bone infection procedure, performed at 10 weeks of age, was adapted from established models ^43^. Briefly, *S. epidermidis* was adjusted to OD_600_ of 5. Titanium pins (0.2 × 0.5 mm cross-section, 4 mm length, L-shape with 1 mm bend) were incubated in the bacterial suspension for 20 min, air-dried for 5 min in a sterile hood, and used within four hours. One pin per batch was placed in 1 mL PBS, sonicated for 3 min, serially diluted, and plated on Tryptic soy agar (TSA, Oxiod, Basel, Switzerland) to determine colony forming units (CFU). Target bacterial load was 10^6-107^ CFU per pin. An inoculated pin was surgically implanted into the right proximal tibia of each mouse.

### Anesthesia, surgery and postoperative care

Tramadol (100 mg/L) was added to drinking water the day before surgery and for 3 days postoperatively. Cages were changed 1–2 days before surgery and placed in a 27° C warming cabinet on the morning of surgery. Anesthesia was induced and maintained using sevoflurane (Baxter, Unterschleissheim, Germany) in oxygen, and buprenorphine (0.1 mg/kg sc, Bupaq, Streuli Tiergesundheit, Uznach, Switzerland) was given as intraoperative analgesia. Warmed Ringer’s lactate (1 mL sc) was administered pre-surgery, and temperature was maintained via heating mat.

The right hindleg was aseptically prepared, and the implant site (2 to 3 mm below the tibial plateau) was identified using the proximal patella as an anatomical landmark. A hole was pre-drilled percutaneously through the proximal tibia from the medial to lateral cortex using a 25-gauge needle. The *S. epidermidis*-inoculated pin was then inserted through the pre-drilled hole, leaving the bent end subcutaneous. If necessary, the puncture was sutured with simple interrupted stitches (Vicryl 5-0, C-3).

Post-surgery, mice were returned to home cages with prior cage mates and kept in the warming cabinet until evening. Recovery food (DietGel^®^ Recovery, ClearH2O, Westbrook, USA) and food soaked in tramadol water were provided on the cage floor. Animals were monitored twice daily for 5 days, daily for 2 days and twice weekly thereafter using a predefined score sheet assessing behavior, lameness, wound healing, Mouse Grimace Scale, respiration, outer appearance (fur), feces and weight.

All mice were euthanized 3 weeks after surgery (13 weeks of age) by intracardial pentobarbital injection (Eskonarkon, Streuli Tiergesundheit, Uznach, Switzerland) under deep sevoflurane anesthesia. Six mice were excluded due to misplaced pins, high blood loss, or a postoperative fracture, resulting in group size variation.

### Bacterial Load Determination

Three weeks post-surgery, bacterial quantification was performed from the right tibia, titanium pin, and surrounding soft tissue. After dissection, samples were collected into sterile PBS (4 mL to soft tissue and bone, and 1 mL to the pin). Soft tissue and tibia were homogenized (Omni Tissue Homogeniser with soft tissue probes, Omni International, Muttenz, Switzerland)), sonicated (3 min) and vortexed. Pins were sonicated (3 min) and vortexed. 1 mL per sample was plated on TSA and incubated for 48 h at 37° C. Colonies were re-streaked on Mannitol Salt Agar (MSA, Oxoid, Basel, Switzerland) for species confirmation. Positive *S. epidermidis* colonies were further plated on TSA with chloramphenicol to verify plasmid-borne antibiotic resistance.

### Sample isolation for flow cytometry

Immediately before euthanasia, 1 mL of blood was collected retrobulbar under anesthesia Samples were lysed twice with 1x red blood cell lysis buffer for 10 min with gentle shaking, followed by quenching in 25 mL of PBS, centrifugation (500 rcf, 5 min), and removal of supernatant.

The right tibia was cleaned, placed in 1 mL of PBS, and transferred to a petri dish containing 2 mL PBS. After removing the bone heads, the bone marrow was flushed with PBS using a 26G needle (Braun, Melsungen, Germany The suspension was disaggregated using a 1000 µL pipette, transferred to a tube with 2 mL of PBS for rinsing, centrifuged (300 rcf, 5 min) and resuspended in 4 mL of PBS.

The spleen was collected and mechanically dissociated over a 70 µm cell strainer (Corning, Berlin, Germany), which was rinsed with 3 mL PBS. The cell suspension was centrifuged (300 rcf, 5 min) and resuspended in 4 mL PBS.

Cell count and viability were assessed for each tissue, and 1×10^6^ cells per sample were plated for staining.

### Antibody surface staining

Cells were distributed in V-bottom 96-well plates, centrifuged (500 xg, 5 min), and resuspended in 40 µL PBS for Live/Dead staining (25 min, 4 °C). After, 160 µL of FACS buffer (PBS containing 1% fetal calf serum (Biochrome, Cambridge, UK)) was added per well, followed by centrifugation (500 xg, 5 min). Extracellular surface staining was performed in 40 μL of Staining Buffer (SB) mixed 1:1 with SB containing Brilliant Violet (BV). Antibodies were applied at the indicated concentrations (Panel innate immune cells and B cells: Table S3; Panel T cells: Table S4, excluding grey markers) for 15 min protected from light. Samples were washed twice with 150 μL SB, centrifuged (500 rcf, 5 min), resuspended in 200 µL FACS buffer, transferred to FACS tubes, and stored at 4° C until same-day acquisition. Alternatively, surface staining was followed by intranuclear staining

### Antibody intranuclear staining

Following surface staining, cells were permeabilized using the Foxp3/ Transcription Factor Staining Buffer Set (eBioscience, San Diego, CA, USA) according to the manufacturer’s protocol. Cells were then resuspended in 40 μL of Foxp3-Buffer containing intracellular staining antibodies (Table S4, markers in grey) and incubated for 30 min at 4 °C in the dark. Subsequently, samples were washed with 160 μL of Foxp3 buffer, resuspended in 200 µL FACS buffer, transferred to FACS tubes, and stored at 4° C until analysis.

### Flow cytometry measurement and analysis

Data were acquired on FACS Aria III using FACSDiva software (Becton Dickinson, San Jose, CA, USA), and analyzed with FlowJo v10 (Treestar, Ashland, OR, US). Gating strategies are detailed in the supplementary information (Figure S12 and S13). Preprocessing was performed in OMIQ (Boston, MA, USA), with arcsinh transformation (coefficient 5), standardized cleanup as described previously ^44^, and quality control using flowCut (Boston, MA, USA). Viable single cells were manually gated.

Total live single cells were downsampled to 10,000 events per group for T cells and 5,000 per subject for innate cells for lineage populations analysis. Dimensionality reduction was performed using unsupervised uniform manifold approximation and projection (UMAP). with parameters: neighbours= 15; minimum distance= 0.4; components= 2; metric= euclidean; embedding initialization= spectral. PhenoGraph clustering was then applied (k = 100).

### *In-vivo* microCT

MicroCT scans of the right proximal tibia were acquired postoperatively (day 0) and at euthanasia (day 21 (VivaCT80, SCANCO Medical AG, Brüttisellen, Switzerland). Anesthesia was induced and maintained with isoflurane during scanning. A 7.49 mm region centered on the surgical pin site, extending from the proximal physis toward the mid-diaphysis, was scanned with a ø21.5 mm field of view and X-ray tube at 70 kVp, 114 µA current with a 0.5 mm Aluminum filter. Scans included 1000 projections over 180° rotation, with a 150 ms integration time per projection (16 min total scan time). Images were acquired at a matrix size of 2066 × 2066 pixels, reconstructed into 720 slices with an isotropic voxel size of 10.4 µm.

### MicroCT Image Processing

Day 0 scans served as a baseline and were rotated to align the tibia along the Y-axis. Day 21 scans were rigidly registered to the baseline scans using bony structures. All datasets were filtered with a 3D Gaussian filter (standard deviation = 1.2, kernel size = 3 × 3 × 3) before evaluation. Bone segmentation was performed on baseline scans using a threshold of 650 mgHA/cm^3^, followed by manual removal of the pin and insertion-induced bone fragments. The resulting segmented data were converted into an intact bone surface model. Five landmarks were manually placed around any observed resorption voids in 3D renderings for day 21 scans. The resorption area was quantified by projecting these landmarks onto the intact bone model and calculating the void area using spline interpolation, excluding the pin volume. All image processing and analysis were performed using custom TCL scripts in Amira (version 2024.2, Thermo Fisher Scientific, Hillsboro, OR, USA).

### Histology

The infected tibia and surrounding soft tissue were fixed in formalin, then decalcified in 12.5% (w/v) EDTA (Roth AG, Arlesheim, Switzerland) and 1.25% (w/v) sodium hydroxide (Sigma-Aldrich). Following decalcification, implants were removed, and the remaining bone and soft tissues were embedded in paraffin (Leica Microsystems). Serial 5 µm sections were prepared using an HM 355 S paraffin microtome (Microm, Thermo Fisher Scientific, Waltham, USA), deparaffinized, and rehydrated through graded ethanol. For each animal, one section containing the pinhole in the bone marrow was stained using Brown & Bren (B&B). Adjacent serial sections were used for multicolor immunofluorescence microscopy.

### Multicolor immunofluorescence staining

Primary antibodies included goat anti-CD3-epsilon (clone M-20, sc-1127, RRID: AB_631128, Santa Cruz Biotechnology) and rabbit anti-*S. aureus* (PA1-7246, RRID:AB_561546, Thermo Fisher Scientific) used at a 1:100 dilution; rabbit-anti Tbet/Tbx21 (MBS248480, MyBiosource), rabbit anti-PD1 (Clone EPR20665, ab214421, RRID:AB_2941806, Abcam), biotin rat anti-Ly6G (clone 1A8, 127604, RRID: AB_1186108, Biolegend), rat anti-mouse Foxp3 (clone FJK-16S, 14-5773-82, RRID:AB_467576, Thermo Fisher Sceintific) and rat anti-mouse Ki67 (HS-398 117, HistoSure) used at a 1:50 dilution.

Secondary antibodies were Alexa Fluor 568-conjugated donkey anti-goat IgG (A-11057, RRID: AB_2534104, Thermo Fisher Scientific) at 1:200 for CD3-epsilon, Alexa Fluor 488-conjugated donkey anti-rabbit IgG (711-546-152, RRID: AB_2340619, Jackson ImmunoResearch Laboratories) at 1:200 for *S. aureus*, Tbet and PD1; and Alexa Fluor 647 donkey anti-rat IgG (712-606-153, RRID: AB_2340696, Jackson ImmunoResearch Laboratories) at 1:200 for Foxp3.

Sections were incubated at 60 °C overnight for deparaffinization, transferred to xylene, and rehydrated through graded ethanol (100%, 96%, 70%) to water. Antigen retrieval was performed in Antigen Unmasking Solution (S1699, Agilent) by boiling for two hours. Nonspecific binding was blocked with 5% donkey serum in Phosphate-Buffered Saline (PBS, MT21040CV, Thermo Fisher Scientific) containing 0.5% Triton X-100 for 40 min in a humidified chamber.

Primary antibodies were applied overnight at 4 °C, followed by PBS washes and two hours incubation with the appropriate secondary antibody at room temperature. Slides were rinsed in PBS for one hour and mounted with Vectashield antifade mounting medium containing DAPI (H-1200-10, Vector Laboratories, Burlingame, USA). Images were acquired using a Zeiss Axioplan 2 microscope equipped with a Hamamatsu camera.

### Data visualization and statistical analysis

Cluster analysis was performed in EdgeR (OMIQ, Boston, MA, USA) to identify significantly different clusters or visualized using heatmaps with Euclidean clustering. Differences in metacluster proportions were assessed by Welch’s t-test on log2-transformed values, followed by Benjamini-Hochberg (BH-)adjustment of p-values (volcano plot).

Other data were visualized using GraphPad Prism 10 (GraphPad Software Inc., La Jolla, CA, US) or R (RStudio: Integrated Development Environment for R. Version 2025.05.1+513, R. Posit, Inc., Boston, MA, US). Statistical significance was determined using either one-way ANOVA with Tukey’s multiple comparisons test or a non-parametric Kruskal-Wallis with Dunn’s multiple comparison test. To address multiple hypothesis testing across cell types while preserving statistical power, only significant post-hoc p-values (p<0.05) plus relevant supplementary cell types were subjected to Benjamini-Hochberg false discovery rate (FDR) correction separately for each figure, reflecting distinct research questions: (1) innate immune response profile, (2) adaptive immune response profile, and (3) cell activation/activity changes. p-values < 0.05 were considered statistically significant and are indicated in the figures. Microscopy images were optimized for brightness and contrast, and scale bars were added using ImageJ v1.53q ^45^ with Fiji ^46^.

## Acknowledgements

The study was supported by SNF (310030 192724) and AO Trauma (AR2021_08 and AR2022_01). JRM is supported by RO1 AI111914. The funder played no role in study design, data collection, analysis and interpretation of data, or the writing of this manuscript.

Virginia Post and Marco Chittò (ARI) are acknowledged for support with *S. epidermidis* transformation and cloning. Iris Keller and Pamela Furlong (ARI) are acknowledged for their support with animal dissection, cell harvesting and staining. Tiziano Schweizer (ARI) is acknowledged for expert advice on antibody panel design. Ursula Menzl (ARI) is acknowledged for flow cytometry machine support. Further, Nadja Von Lanthen and Corinne Bischofberger (ARI) are acknowledged for support with the dissection of the mice. Nico Valerio Giger (ARI) is acknowledged for his help with programming in R. Daniel Arens and Claudia Zindl (ARI) are acknowledged for support with animal handling and surgery assistance. Antonin Weckel and Tiffany Scharschmidt are acknowledged for the initial advice to set up the study and provide the plasmid. Further we would like to acknowledge the animal caretakers and the mice.

## Data Availability

The datasets generated and analysed during the current study are available in the Zenodo repository, “https://zenodo.org/records/18172448”.

## Contributions

PF and EMAK contributed equally to data collection, data curation, data visualization, manuscript writing and manuscript revision. LG and JTD contributed both to data collection, manuscript writing, and manuscript revision. JRM contributed to histology data curation, data visualization, data interpretation, manuscript revision and funding. AH contributed to data collection, manuscript writing and manuscript revision. PK contributed to data visualization. SZ contributed to manuscript revision. ECDJ contributed to data interpretation and manuscript revision. GM contributed to data interpretation, manuscript revision and funding. TFM contributed to supervision, funding, and manuscript revision.

## References

1 Brown, M. M. & Horswill, A. R. Staphylococcus epidermidis-Skin friend or foe? PLoS.Pathog 16, e1009026 (2020). 10.1371/journal.ppat.1009026

2 Scharschmidt, T. C.. et al.¡ A Wave of Regulatory T Cells into Neonatal Skin Mediates Tolerance to Commensal Microbes. Immunity 43, 1011–1021 (2015). 10.1016/j.immuni.2015.10.016

3 Parlet, C. P., Brown, M. M. & Horswill, A. R. Commensal Staphylococci Influence Staphylococcus aureus Skin Colonization and Disease. Trends.Microbiol 27, 497–507 (2019). 10.1016/j.tim.2019.01.008

4 Leshem, A., Liwinski, T. & Elinav, E. Immune-Microbiota Interplay and Colonization Resistance in Infection. Mol.Cell 78, 597–613 (2020). 10.1016/j.molcel.2020.03.001

5 Severn, M. M. & Horswill, A. R. Staphylococcus epidermidis and its dual lifestyle in skin health and infection. Nat.Rev.Microbiol 21, 97–111 (2023). 10.1038/s41579-022-00780-3

6 Weckel, A.. et al.¡ Long-term tolerance to skin commensals is established neonatally through a specialized dendritic cell subgroup. Immunity 56, 1239-1254.e1237 (2023). 10.1016/j.immuni.2023.03.008

7 Volz, T.. et al.¡ Induction of IL-10-balanced immune profiles following exposure to LTA from Staphylococcus epidermidis. Exp. Dermatol 27, 318–326 (2018). 10.1111/exd.13540

8 Belkaid, Y., Bouladoux, N. & Hand, T. W. Effector and memory T cell responses to commensal bacteria. Trends.Immunol 34, 299–306 (2013). 10.1016/j.it.2013.03.003

9 Sakaguchi, S.. et al.¡ Regulatory T Cells and Human Disease. Annu. Rev.Immunol 38, 541–566 (2020). 10.1146/annurev-immunol-042718-041717

10 Naik, S.. et al.¡ Commensal-dendritic-cell interaction specifies a unique protective skin immune signature. Nature 520, 104–108 (2015). 10.1038/nature14052

11 Leech, J. M.. et al.¡ Toxin-Triggered Interleukin-1 Receptor Signaling Enables Early-Life Discrimination of Pathogenic versus Commensal Skin Bacteria. Cell.Host.Microbe 26, 795-809.e795 (2019). 10.1016/j.chom.2019.10.007

12 Scharschmidt, T. C. Establishing Tolerance to Commensal Skin Bacteria: Timing Is Everything. Dermatol.Clin 35, 1–9 (2017). 10.1016/j.det.2016.07.007

13 Schäfer, P.. et al.¡ Prolonged bacterial culture to identify late periprosthetic joint infection: a promising strategy. Clin.Infect.Dis 47, 1403–1409 (2008). 10.1086/592973

14 Moriarty, T. F.. et al.¡ Orthopaedic device-related infection: current and future interventions for improved prevention and treatment. EFORT.Open.Rev 1, 89–99 (2016). 10.1302/2058-5241.1.000037

15 Pius, A. K.. et al.¡ Effects of Aging on Osteosynthesis at Bone-Implant Interfaces. Biomolecules 14 (2023). 10.3390/biom14010052

16 Nguyen, T. H., Park, M. D. & Otto, M. Host Response to Staphylococcus epidermidis Colonization and Infections. Front. Cell.Infect.Microbiol 7, 90 (2017). 10.3389/fcimb.2017.00090

17 Schilcher, K. & Horswill, A. R. Staphylococcal Biofilm Development: Structure, Regulation, and Treatment Strategies. Microbiol.Mol.Biol.Rev 84 (2020). 10.1128/mmbr.00026-19

18 Mah, T. F. & O’Toole, G. A. Mechanisms of biofilm resistance to antimicrobial agents. Trends.Microbiol 9, 34–39 (2001). 10.1016/s0966-842x(00)01913-2

19 de Vor, L., Rooijakkers, S. H. M. & van Strijp, J. A. G. Staphylococci evade the innate immune response by disarming neutrophils and forming biofilms. FEBS.Lett 594, 2556–2569 (2020). 10.1002/1873-3468.13767

20 Lee, J. Y. H.. et al.¡ Global spread of three multidrug-resistant lineages of Staphylococcus epidermidis. Nat.Microbiol 3, 1175–1185 (2018). 10.1038/s41564-018-0230-7

21 Salgueiro, V. C.. et al.¡ High rate of neonates colonized by methicillin-resistant Staphylococcus species in an Intensive Care Unit. J.Infect.Dev.Ctries 13, 810–816 (2019). 10.3855/jidc.11241

22 Perera Molligoda Arachchige, A.S. Human NK cells: From development to effector functions. Innate.Immun 27, 212–229 (2021). 10.1177/17534259211001512

23 Sottnik, J. L., U’Ren, L. W., Thamm, D. H., Withrow, S. J. & Dow, S. W. Chronic bacterial osteomyelitis suppression of tumor growth requires innate immune responses. Cancer.Immunol. Immunother 59, 367–378 (2010). 10.1007/s00262-009-0755-y

24 Zhao, Y., Liu, Q., Zhao, J. & Song, D. The roles of natural killer cells in bone and arthritic disease: a narrative review. Immunol.Med 48, 281–294 (2025). 10.1080/25785826.2025.2506260

25 Nimmana, B. K. & Savaliya, V. in StatPearls (StatPearls Publishing Copyright © 2025, StatPearls Publishing LLC., 2025).

26 Jaeger, B. N.. et al.¡ Neutrophil depletion impairs natural killer cell maturation, function, and homeostasis. J.Exp.Med 209, 565–580 (2012). 10.1084/jem.20111908

27 Sabaté-Brescó, M.. et al.¡ Fracture biomechanics influence local and systemic immune responses in a murine fracture-related infection model. Biol.Open 10 (2021). 10.1242/bio.057315

28 O’Keeffe, M., Mok, W. H. & Radford, K. J. Human dendritic cell subsets and function in health and disease. Cell.Mol.Life.Sci 72, 4309–4325 (2015). 10.1007/s00018-015-2005-0

29 Wang, G.. et al.¡ “Default” generation of neonatal regulatory T cells. J.Immunol 185, 71–78 (2010). 10.4049/jimmunol.0903806

30 Nutsch, K.. et al.¡ Rapid and Efficient Generation of Regulatory T Cells to Commensal Antigens in the Periphery. Cell.Rep 17, 206–220 (2016). 10.1016/j.celrep.2016.08.092

31 García-Ojeda, M. E., Dejbakhsh-Jones, S., Weissman, I. L. & Strober, S. An alternate pathway for T cell development supported by the bone marrow microenvironment: recapitulation of thymic maturation. J.Exp.Med 187, 1813–1823 (1998). 10.1084/jem.187.11.1813

32 Surendar, J.. et al.¡ Osteomyelitis is associated with increased anti-inflammatory response and immune exhaustion. Front. Immunol 15, 1396592 (2024). 10.3389/fimmu.2024.1396592

33 Sedney, C. J. & Harvill, E. T. The Neonatal Immune System and Respiratory Pathogens. Microorganisms 11 (2023). 10.3390/microorganisms11061597

34 Odorizzi, P. M., Pauken, K. E., Paley, M. A., Sharpe, A. & Wherry, E. J. Genetic absence of PD-1 promotes accumulation of terminally differentiated exhausted CD8+ T cells. J.Exp.Med 212, 1125–1137 (2015). 10.1084/jem.20142237

35 Jubel, J. M., Barbati, Z. R., Burger, C., Wirtz, D. C. & Schildberg, F. A. The Role of PD-1 in Acute and Chronic Infection. Front. Immunol 11, 487 (2020). 10.3389/fimmu.2020.00487

36 Wei, F.. et al.¡ Strength of PD-1 signaling differentially affects T-cell effector functions. Proceedings.of.the.National.Academy.of. Sciences 110, E2480–E2489 (2013). doi:10.1073/pnas.1305394110

37 Guarch-Pérez, C., Riool, M. & Zaat, S. A. Current osteomyelitis mouse models, a systematic review. Eur.Cell.Mater 42, 334–374 (2021). 10.22203/eCM.v042a22

38 Sabaté Brescó, M.. et al.¡ Pathogenic Mechanisms and Host Interactions in Staphylococcus epidermidis Device-Related Infection. Front.Microbiol 8, 1401 (2017). 10.3389/fmicb.2017.01401

39 Cole, L. E.. et al.¡ Limitations of Murine Models for Assessment of Antibody-Mediated Therapies or Vaccine Candidates against Staphylococcus epidermidis Bloodstream Infection. Infect. Immun 84, 1143–1149 (2016). 10.1128/iai.01472-15

40 Gantenbein, F.. et al.¡ Protocol for a systematic review of good surgical practice guidelines for experimental rodent surgery. BMJ. Open.Sci 6, e100280 (2022). 10.1136/bmjos-2022-100280

41 Daoust, L.. et al.¡ Gnotobiotic mice housing conditions critically influence the phenotype associated with transfer of faecal microbiota in a context of obesity. Gut 72, 896–905 (2023). 10.1136/gutjnl-2021-326475

42 Löfblom, J., Kronqvist, N., Uhlén, M., Ståhl, S. & Wernérus, H. Optimization of electroporation-mediated transformation: Staphylococcus carnosus as model organism. J.Appl.Microbiol 102, 736–747 (2007). 10.1111/j.1365-2672.2006.03127.x

43 Li, D.. et al.¡ Quantitative mouse model of implant-associated osteomyelitis and the kinetics of microbial growth, osteolysis, and humoral immunity. J.Orthop.Res 26, 96–105 (2008). 10.1002/jor.20452

44 Bagwell, C. B.. et al.¡ Automated Data Cleanup for Mass Cytometry. Cytometry.A 97, 184–198 (2020). 10.1002/cyto.a.23926

45 Schneider, C. A., Rasband, W. S. & Eliceiri, K. W. NIH Image to ImageJ: 25 years of image analysis. Nat.Methods 9, 671–675 (2012). 10.1038/nmeth.2089

46 Schindelin, J.. et al.¡ Fiji: an open-source platform for biological-image analysis. Nat.Methods 9, 676–682 (2012). 10.1038/nmeth.2019

