## Supplementary Material for "Beyond the skin barrier: commensal *S. epidermidis* imprint systemic immunity to invasive biofilm infection"

### Supplementary data

Table S1 shows the significantly different clusters and the corresponding immune cell types in bone marrow for Figure 1. A) Comparing non-colonized (Non) with neonatal-colonized (NeoC) mice, upregulated clusters in NeoC in white and upregulated in Non in grey. B) Comparing Non with adult-colonized (AdC) mice, upregulated clusters in AdC in white and upregulated in Non in grey. C) Comparing NeoC with AdC mice, upregulated clusters in AdC in white and upregulated in NeoC in grey.

*Table S1. Significantly different clusters and corresponding Immune cell types*

**A** **Non vs NeoC**

| Cell type | Marker | Cluster(s) |
|---|---|---|
| NK cells | NK1.1$^+$ | C22 ($p$<0.001), C01 ($p$<0.001), C08 ($p$<0.001), C21 ($p$<0.001), C17 ($p$<0.001), C02 ($p$<0.001), C07 ($p$<0.001) |
| B cells | CD19$^+$, CD20$^+$ | C06 ($p$<0.001), C11 ($p$<0.001), C05 ($p$=0.003), C16 ($p$=0.009) |
| Monocytes | Ly6c$^+$ | C34 ($p$<0.001), C25 ($p$=0.032) |
| T cells | CD3$^+$ | C18 ($p$<0.001), C24 ($p$<0.001) |
| DCs | CD80$^+$ | C36 ($p$=0.005) |
| Macrophages | CD11b$^+$ | C28 ($p$=0.033) |
| NK cells | NK1.1$^+$ | C20 ($p$<0.001), C04 ($p$<0.001), C29 ($p$=0.003) |
| B cells | CD19$^+$, CD20$^+$ | C12 ($p$<0.001), C27 ($p$<0.001), C02 ($p$<0.001) C19 ($p$=0.003) |
| T cells | CD3$^+$ | C10 ($p$=0.009) |
| Macrophages | CD11b$^+$ | C32 ($p$=0.022) |
| Macrophages | F4/80$^+$ | C38 ($p$=0.041) |

**B** **Non vs AdC**

| Cell type | Marker | Cluster(s) |
|---|---|---|
| NK cells | NK1.1$^+$ | C01 ($p$<0.001), C21 ($p$<0.001), C17 ($p$<0.001), C08 ($p$=0.002), C07 ($p$=0.003), C25 ($p$=0.017), C03 ($p$=0.020) |
| B cells | CD19$^+$, CD20$^+$ | C11 ($p$<0.001), C15 ($p$=0.002), C05 ($p$=0.010), C16 ($p$=0.018), C27 ($p$=0.039) |
| T cells | CD3$^+$ | C24 ($p$<0.001), C13 ($p$=0.019), C33 ($p$=0.029) |
| Monocytes | Ly6c$^+$ | C35 ($p$=0.047) |
| Macrophages | CD11b$^+$ | C28 ($p$=0.009) |
| B cells | CD19$^+$, CD20$^+$ | C04 ($p$<0.001) |
| NK cells | NK1.1$^+$ | C20 ($p$=0.002), C29 ($p$=0.008) |
| Macrophages | CD11b$^+$ | C32 ($p$=0.029) |

**C** **NeoC vs AdC**

| Cell type | Marker | Cluster(s) |
|---|---|---|
| T cells | CD3$^+$ | C13 ($p$<0.001) |
| B cells | CD19$^+$, CD20$^+$ | C27 ($p$<0.001), C02 ($p$<0.001), C12 ($p$<0.001), C19 ($p$=0.016) |
| Monocytes | Ly6c$^+$ | C35 ($p$=0.020) |
| NK cells | NK1.1$^+$ | C22 ($p$=0.004) |
| T cells | CD3$^+$ | C18 ($p$=0.007) |
| B cells | CD19$^+$, CD20$^+$ | C06 ($p$=0.008) |
| Macrophages | CD11b$^+$ | C34 ($p$=0.009) |

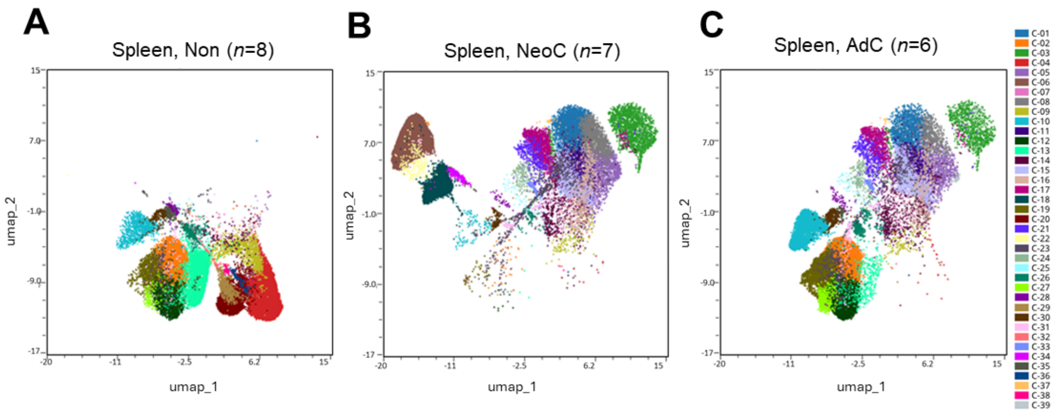


**D** **Non vs NeoC**

| Cell type | Marker | Cluster(s) |
|---|---|---|
| NK cells | NK1.1$^+$ | C08 ($p<0.001$), C01 ($p<0.001$), C17 ($p<0.001$), C21 ($p<0.001$), C22 ($p=0.009$), C03 ($p=0.00013$) |
| B cells | CD19$^+$, CD20$^+$ | C06 ($p<0.001$), C11 ($p<0.001$), C15 ($p<0.001$), C05 ($p=0.005$), C16 ($p=0.041$), C37 ($p=0.037$) |
| Monocytes | Ly6c$^+$ | C34 ($p=0.002$) |
| T cells | CD3$^+$ | C18 ($p=0.001$), C33 ($p=0.003$), C24 ($p=0.009$) |
| NK cells | NK1.1$^+$ | C20 ($p<0.001$), C04 ($p<0.001$), C29 ($p=0.004$) |
| B cells | CD19$^+$, CD20$^+$ | C12 ($p=0.003$), C27 ($p=0.005$), C02 ($p=0.008$) |
| T cells | CD3$^+$ | C13 ($p<0.001$), C38 ($p=0.006$) |
| Monocytes | Ly6c$^+$ | C35 ($p=0.040$) |
| DCs | CD80$^+$ | C36 ($p=0.010$) |

**E** **Non vs AdC**

| Cell type | Marker | Cluster(s) |
|---|---|---|
| NK cells | NK1.1$^+$ | C01 ($p=0.003$), C17 ($p=0.003$), C08 ($p=0.004$), C21 ($p=0.008$), C03 ($p=0.008$) |
| B cells | CD19$^+$, CD20$^+$ | C11 ($p=0.003$), C15 ($p=0.003$), C05 ($p=0.016$), C16 ($p=0.046$), C39 ($p=0.032$) |
| T cells | CD3$^+$ | C33 ($p=0.003$), C24 ($p=0.012$) |
| B cells | CD19$^+$, CD20$^+$ | C04 ($p=0.004$) |
| NK cells | NK1.1$^+$ | C20 ($p=0.006$), C29 ($p=0.018$) |
| Macrophages | CD11b$^+$ | C32 ($p=0.045$) |
| DCs | CD80$^+$ | C36 ($p=0.027$) |
| T cells | CD3$^+$ | C38 ($p=0.011$) |

**F** **NeoC vs AdC**

| Cell type | Marker | Cluster(s) |
|---|---|---|
| T cells | CD3$^+$ | C13 ($p<0.001$), C30 ($p=0.034$) |
| B cells | CD19$^+$, CD20$^+$ | C27 ($p<0.001$), C02 ($p=0.001$), C12 ($p<0.001$), C19 ($p=0.033$) |
| NK cells | NK1.1$^+$ | C22 ($p=0.019$) |
| T cells | CD3$^+$ | C10 ($p=0.003$), C18 ($p=0.008$) |
| B cells | CD19$^+$, CD20$^+$ | C06 ($p=0.007$) |
| Macrophages | CD11b$^+$ | C34 ($p=0.015$) |

**Figure S1. Immune cells compared between colonization groups in the blood.** Uniform Manifold Approximation and Projection (UMAP) representative performed based on marker expression profiles of all significantly different clusters for **A)** non-colonized (Non; n=8), **B)** neonatal-colonized (NeoC; n=8), and **C)** adult-colonized (AdC; n=6) mice. **D)** Significantly different clusters and corresponding cell type comparing Non with NeoC, upregulated clusters in NeoC in white and upregulated in Non in grey. **E)** Comparing Non with AdC, upregulated clusters in AdC in white and upregulated in Non in grey. **F)** Comparing NeoC with AdC mice, upregulated clusters in AdC in white and upregulated in NeoC in grey.

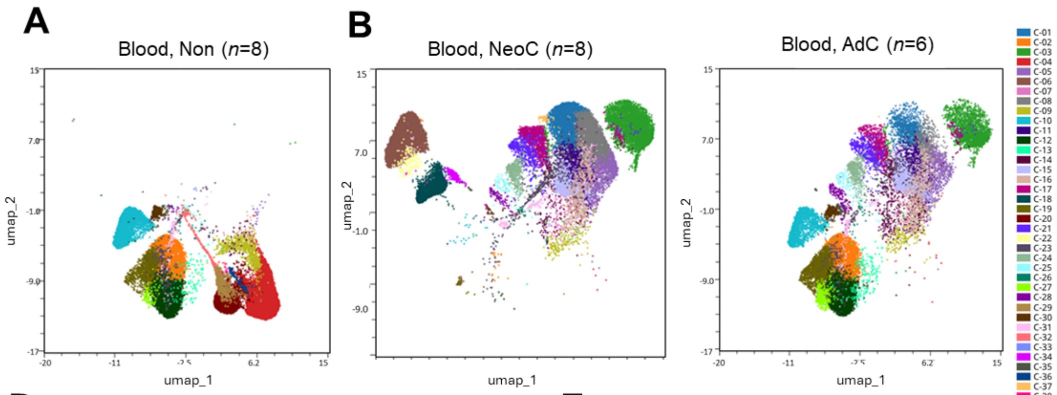


**D** **Non vs NeoC**

| Cell type | Marker | Cluster(s) |
|---|---|---|
| NK cells | NK1.1⁺ | C01 ($p$<0.001), C08 ($p$<0.001), C21 ($p$<0.001), C17 ($p$<0.001), C03 ($p$<0.001), C22 ($p$=0.002), C05 ($p$=0.003), C07 ($p$=0.012) |
| B cells | CD19⁺, CD20⁺ | C06 ($p$<0.001), C11 ($p$<0.001), C15 ($p$<0.001), C37 ($p$=0.009), C16 ($p$=0.037) |
| Monocytes | Ly6c⁺ | C34 ($p$<0.001) |
| T cells | CD3⁺ | C18 ($p$<0.001) |
| | | C24 ($p$<0.001) |
| NK cells | NK1.1⁺ | C20 ($p$<0.001), C04 ($p$<0.001), C29 ($p$<0.001) |
| B cells | CD19⁺, CD20⁺ | C12 ($p$<0.001), C02 ($p$<0.001), C27 ($p$=0.010), C19 ($p$=0.044) |
| T cells | CD3⁺ | C10 ($p$=0.004), C13 ($p$=0.017) |
| Macrophages | CD11b⁺ | C32 ($p$=0.004) |
| DCs | CD80⁺ | C36 ($p$=0.002) |

**E** **Non vs AdC**

| Cell type | Marker | Cluster(s) |
|---|---|---|
| NK cells | NK1.1⁺ | C03 ($p$<0.001), C21 ($p$<0.001), C17 ($p$=0.003), C08 ($p$=0.004), C05 ($p$=0.004), C21 ($p$=0.006), C07 ($p$=0.023) |
| B cells | CD19⁺, CD20⁺ | C01 ($p$=0.001), C15 ($p$=0.002), C11 ($p$=0.003), C14 ($p$=0.029), C16 ($p$=0.030) |
| T cells | CD3⁺ | C24 ($p$<0.001) |
| Macrophages | CD11b⁺ | C28 ($p$=0.008) |
| B cells | CD19⁺, CD20⁺ | C04 ($p$=0.002) |
| NK cells | NK1.1⁺ | C20 ($p$=0.008), C29 ($p$=0.012) |
| Macrophages | CD11b⁺ | C32 ($p$=0.014) |
| DCs | CD80⁺ | C36 ($p$=0.008) |

**F** **NeoC vs AdC**

| Cell type | Marker | Cluster(s) |
|---|---|---|
| T cells | CD3⁺ | C13 ($p$<0.001), C10 ($p$=0.003), C30 ($p$=0.034) |
| B cells | CD19⁺, CD20⁺ | C27 ($p$<0.001), C12 ($p$<0.001), C02 ($p$=0.001), C19 ($p$=0.032) |
| NK cells | NK1.1⁺ | C22 ($p$=0.019) |
| T cells | CD3⁺ | C18 ($p$=0.008) |
| B cells | CD19⁺, CD20⁺ | C06 ($p$=0.007) |
| Macrophages | CD11b⁺ | C34 ($p$=0.015) |

**Figure S2. Splenic immune cells compared between colonization groups.** Uniform Manifold Approximation and Projection (UMAP) representative performed based on marker expression profiles of all significantly different clusters for **A)** non-colonized (Non; n=8), **B)** neonatal-colonized (NeoC; n=8), and **C)** adult-colonized (AdC; n=6) mice. **D)** Significantly different clusters and corresponding cell type comparing Non with NeoC, upregulated clusters in NeoC in white and upregulated in Non in grey. **E)** Comparing Non with AdC, upregulated clusters in AdC in white and upregulated in Non in grey. **F)** Comparing NeoC with AdC mice, upregulated clusters in AdC in white and upregulated in NeoC in grey.

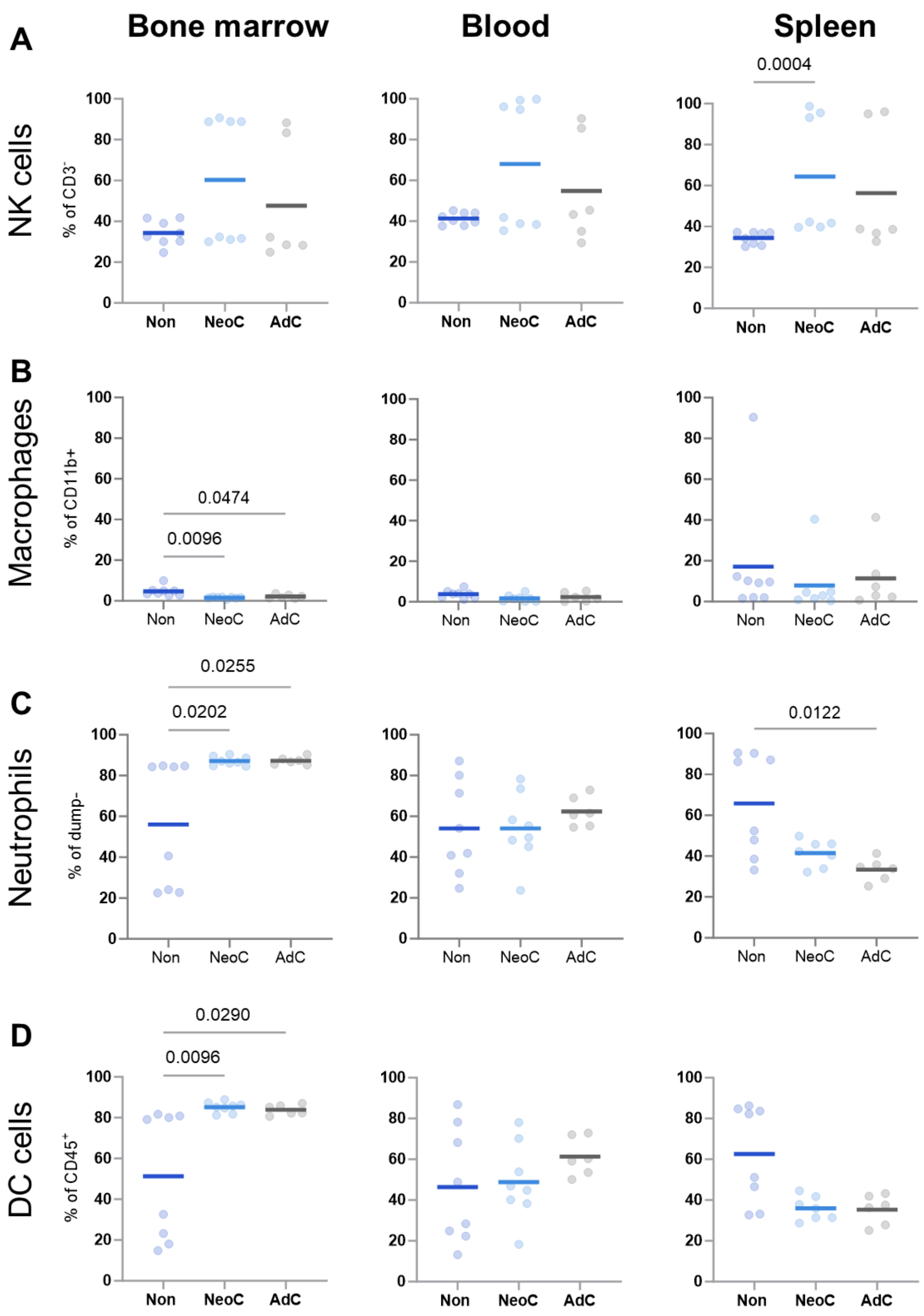


**Figure S3. Innate immune cell abundance in bone marrow, blood and spleen. A)** Natural Killer (NK) cells, **B)** macrophages, **C)** neutrophils and, **D)** Dendritic Cells (DCs) in bone marrow, blood and spleen between non-colonized (Non: *n=8*), neonatal-colonized (NeoC: *n=8*) and adult-colonized (AdC: *n=6*) groups from individual mice shown as dots with mean as line. One sample was lost for NeoC NK cells in the spleen (*n=7*). Statistical analyses performed using one-way ANOVA with Tukey's multiple comparisons test or non-parametric with Kruskal-Wallis test. Only significant p-values shown.

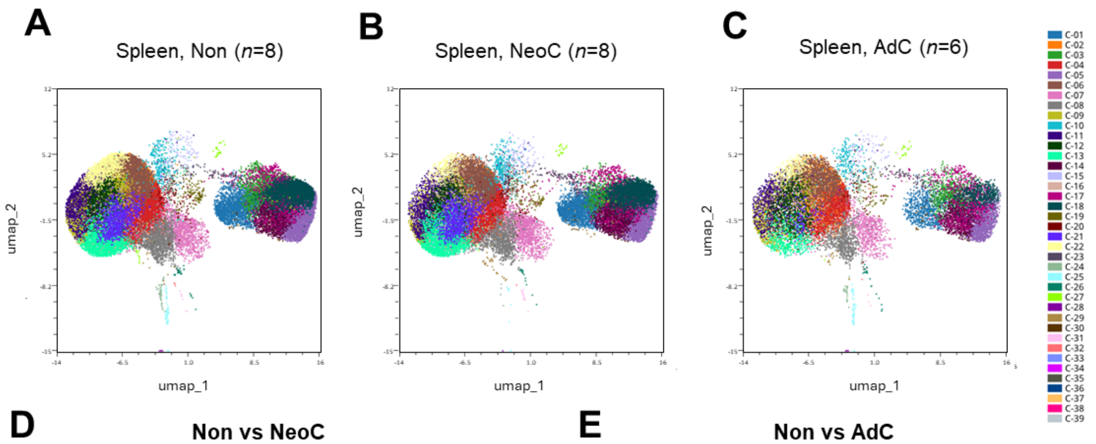


**D** Non vs NeoC

| Cell type | Marker | Cluster(s) |
|---|---|---|
| Th2 | CD4$^{+}$, IL-4$^{+}$ | C04 ($p$<0.001),<br>C13 ($p$<0.001),<br>C21 ($p$<0.001) |
| Effector cells | CD4$^{+}$,CD44$^{+}$ | C22 ($p$<0.001),<br>C26 ($p$<0.001),<br>C09 ($p$= 0.012),<br>C08 ($p$=0.043) |
| Th2 | CD4$^{+}$, IL-4$^{+}$ | C19 ($p$<0.001),<br>C25 ($p$<0.001),<br>C27 ($p$<0.001),<br>C24 ($p$=0.004),<br>C10 ($p$=0.009),<br>C07 ($p$=0.029),<br>C15 ($p$=0.044) |
| Treg | CD4$^{+}$,FoxP3$^{+}$ | C28 ($p$<0.001) |
| CD8$^{+}$ T cells | CD4$^{+}$,CD8$^{+}$ | C22 ($p$<0.001),<br>C03 ($p$=0.027),<br>C31 ($p$=0.048) |

**E** Non vs AdC

| Cell type | Marker | Cluster(s) |
|---|---|---|
| Th2 | CD4$^{+}$, IL-4$^{+}$ | C04 ($p$<0.001),<br>C13 ($p$<0.001),<br>C21 ($p$=0.007) |
| Treg | CD4$^{+}$,FoxP3$^{+}$ | C28 ($p$=0.048) |
| Th2 | CD4$^{+}$, IL-4$^{+}$ | C27 ($p$=0.009) |

**F** NeoC vs AdC

| Cell type | Marker | Cluster(s) |
|---|---|---|
| Th2 | CD4$^{+}$, IL-4$^{+}$ | C19 ($p$<0.001) |
| Treg | CD4$^{+}$,FoxP3$^{+}$ | C28 ($p$=0.001) |
| CD8$^{+}$ T cells | CD4$^{+}$,CD8$^{+}$ | C22 ($p$=0.002) |
| Th2 | CD4$^{+}$, IL-4$^{+}$ | C25 ($p$<0.001),<br>C24 ($p$=0.016),<br>C19 ($p$=0.002),<br>C09 ($p$=0.004),<br>C26 ($p$=0.007),<br>C02 ($p$=0.016) |
| Effector cells | CD4$^{+}$,CD44$^{+}$ | C08 ($p$=0.017) |
| CD8$^{+}$ T cells | CD4$^{+}$,CD8$^{+}$ | C18 ($p$=0.016) |
| Th2 | CD4$^{+}$, IL-4$^{+}$ | C21 ($p$=0.013) |

**Figure S4. T cells compared between colonization groups in the spleen.** Uniform Manifold Approximation and Projection (UMAP) representative performed based on marker expression profiles of all significantly different clusters for **A)** non-colonized (Non; n=8), **B)** neonatal-colonized (NeoC; n=8), and **C)** adult-colonized (AdC; n=6) mice. **D)** Significantly different clusters and corresponding cell type comparing Non with NeoC, upregulated clusters in NeoC in white and upregulated in Non in grey. **E)** Comparing Non with AdC, upregulated clusters in AdC in white and upregulated in Non in grey. **F)** Comparing NeoC with AdC mice, upregulated clusters in AdC in white and upregulated in NeoC in grey.

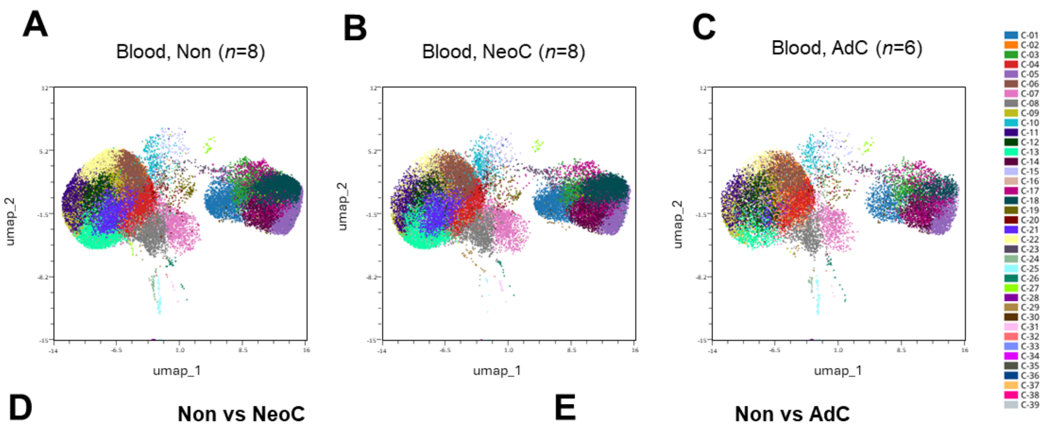


**D** **Non vs NeoC**

| Cell type | Marker | Cluster(s) |
|---|---|---|
| Th2 | $CD4^+$, IL-$4^+$ | C29 ($p$=0.006) |
| Treg | $CD4^+$,$FoxP3^+$ | C28 ($p$<0.001) |

**E** **Non vs AdC**

| Cell type | Marker | Cluster(s) |
|---|---|---|
| $CD8^+$ T cells | $CD4^+$,$CD8^+$ | C23 ($p$=0.027) |
| Treg | $CD4^+$,$FoxP3^+$ | C28($p$=0.003) |
| Th2 | $CD4^+$, IL-$4^+$ | C24 ($p$=0.009),<br>C29 ($p$=0.009),<br>C25 ($p$=0.010) |

**F** **NeoC vs AdC**

| Cell type | Marker | Cluster(s) |
|---|---|---|
| Treg | $CD4^+$,$FoxP3^+$ | C28 ($p$<0.001) |
| Th2 | $CD4^+$, IL-$4^+$ | C25 ($p$<0.001)<br>C24 ($p$=0.002)<br>C26 ($p$=0.031) |

**Figure S5. T cells compared between colonization groups in the blood.** Uniform Manifold Approximation and Projection (UMAP) representative performed based on marker expression profiles of all significantly different clusters for **A)** non-colonized (Non; n=8), **B)** neonatal-colonized (NeoC; n=8), and **C)** adult-colonized (AdC; n=6) mice. **D)** Significantly different clusters and corresponding cell type comparing Non with NeoC, upregulated clusters in NeoC in white and upregulated in Non in grey. **E)** Comparing Non with AdC, upregulated clusters in AdC in white and upregulated in Non in grey. **F)** Comparing NeoC with AdC mice, upregulated clusters in AdC in white and upregulated in NeoC in grey.

Table S2 shows the significant different clusters and the corresponding T cell types in the bone marrow from Figure 2 comparing **A)** non-colonized (Non; n=8) with neonatal-colonized (NeoC; n=8), upregulated clusters in NeoC in white and upregulated in Non in grey. **B)** Comparing Non with adult-colonized (AdC; n=6), upregulated clusters in AdC in white and upregulated in Non in grey. **C)** Comparing NeoC with AdC mice, upregulated clusters in AdC in white and upregulated in NeoC in grey.

*Table S2. Significantly different clusters and corresponding T cell types*

**A** **Non vs NeoC**

| Cell type | Marker | Cluster(s) |
|---|---|---|
| Effector cells | $CD4^+$,$CD44^+$ | C09 ($p$=0.007), C12 ($p$=0.015) |
| Th2 | $CD4^+$, $IL\text{-}4^+$ | C04 ($p$<0.001), C05 ($p$=0.015), C13 ($p$=0.019), C20 ($p$=0.038) |
| Th2 | $CD4^+$, $IL\text{-}4^+$ | C04 ($p$<0.001), C11 ($p$=0.02), C15 ($p$=0.028) |

**B** **NeoC vs AdC**

| Cell type | Marker | Cluster |
|---|---|---|
| Treg | $CD4^+$, $FOX\text{-}P3^+$ | C18 ($p$<0.001) |
| Th2 | $CD4^+$, $IL\text{-}4^+$ | C14 ($p$<0.001), C17 ($p$<0.001), C15 ($p$<0.001) |
| Th2 | $CD4^+$, $IL\text{-}4^+$ | C11 ($p$=0.012) |

**C** **Non vs AdC**

| Cell type | Marker | Cluster(s) |
|---|---|---|
| Treg | $CD4^+$, $FOX\text{-}P3^+$ | C18 ($p$=0.006) |
| Effector cells | $CD4^+$,$CD44^+$ | C09 ($p$=0.033), C12 ($p$=0.004) |
| Th2 | $CD4^+$, $IL\text{-}4^+$ | C14 ($p$=0.001), C17 ($p$=0.001), C15 ($p$=0.004), C13 ($p$=0.014), C19 ($p$=0.038) |
| Th2 | $CD4^+$, $IL\text{-}4^+$ | C04 ($p$<0.001), C11 ($p$=0.001) |

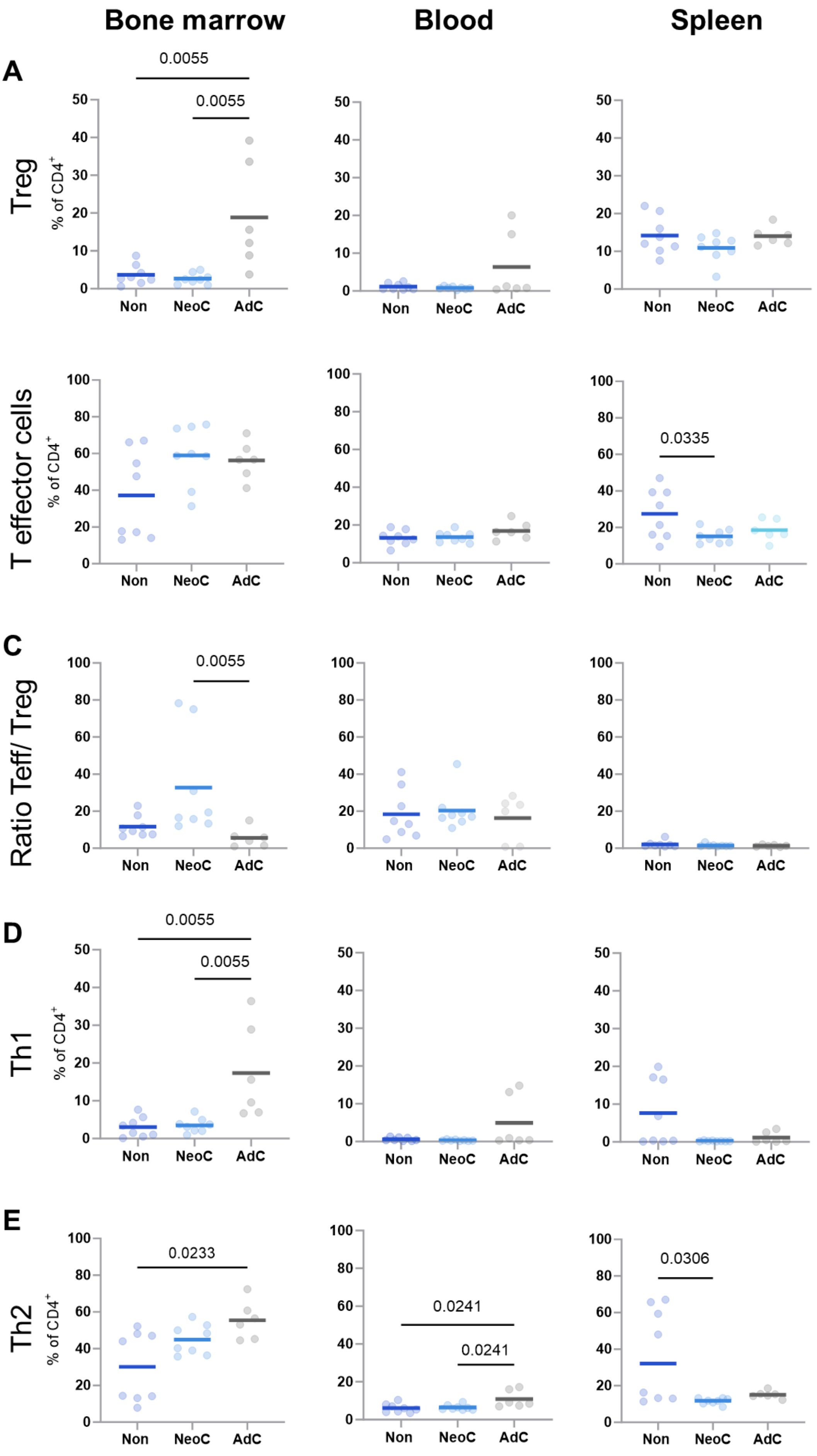
Bone marrow
Blood
Spleen
A
Treg
% of CD4+
0.0055
0.0055
Non
NeoC
AdC
T effector cells
0.0335
C
Ratio Teff/ Treg
0.0055
D
Th1
0.0055
0.0055
E
Th2
0.0233
0.0241
0.0241
0.0306

**Figure S6. T cell subpopulation abundance in bone marrow, blood and spleen. A)** T regulatory (Treg), **B)** T effector (Teff), **C)** the ration between Teff to Treg, **D)** T helper 1 (Th1) and, **E)** T helper 2 (Th2) cells in bone marrow, blood and spleen between non-colonized (Non: *n=8*), neonatal-colonized (NeoC: *n=8*) and adult-colonized (AdC: *n=6*) groups from individual mice shown as dots with mean as line. Statistical analyses performed using one-way ANOVA with Tukey's multiple comparisons test or non-parametric with Kruskal-Wallis test. Only significant p-values shown.

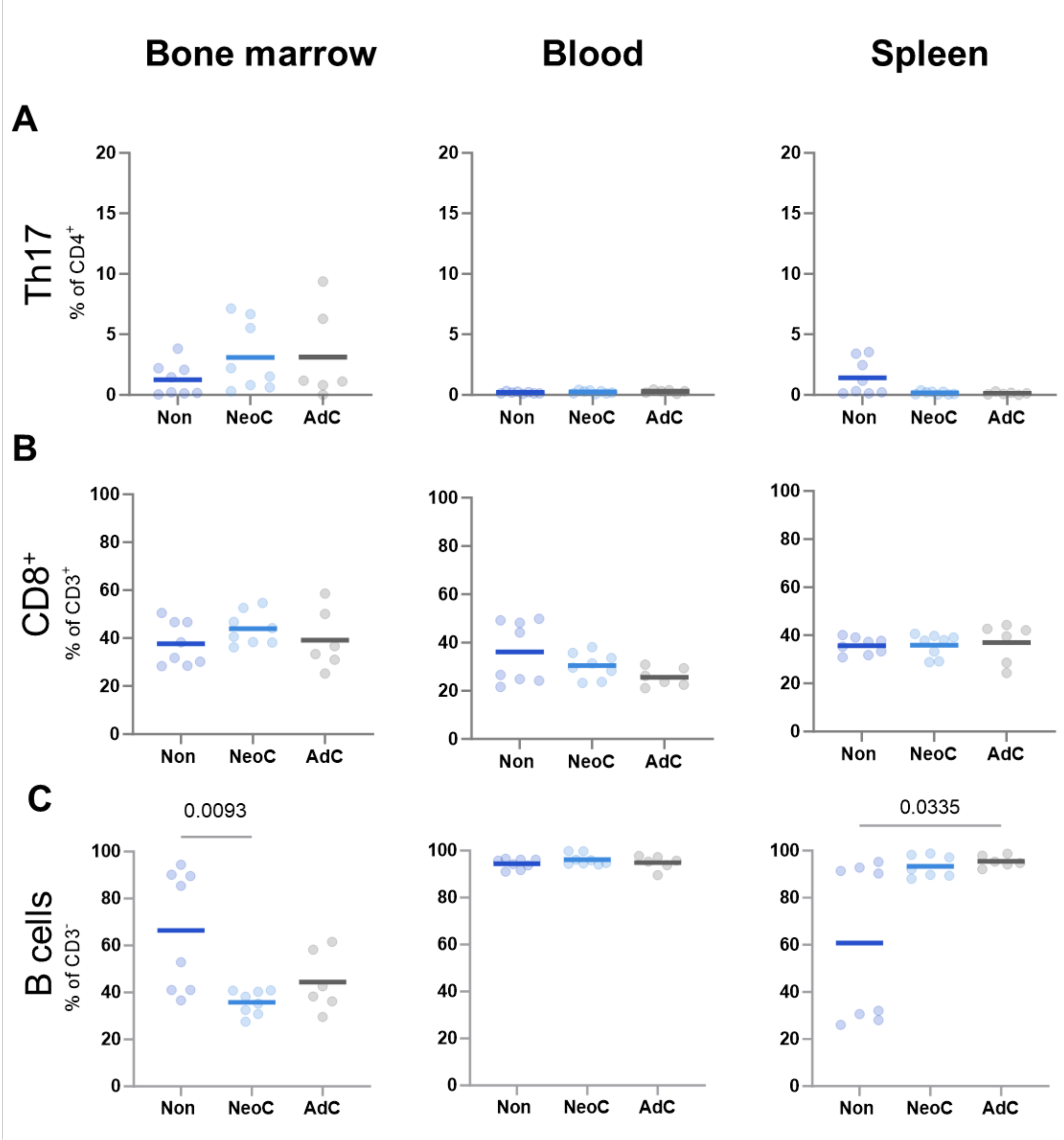


**Figure S7. Th17, $CD8^+$ T cells and B cells in bone marrow, blood and spleen. A)** T helper 17 (Th17), **B)** $CD8^+$ T cells, and **C)** B cells in bone marrow, blood and spleen between non-colonized (Non: n=8), neonatal-colonized (NeoC: n=8) and adult-colonized (AdC: n=6) groups from individual mice shown as dots with mean as line. Statistical analyses performed using one-way ANOVA with Tukey's multiple comparisons test or non-parametric with Kruskal-Wallis test. Only significant p-values shown.

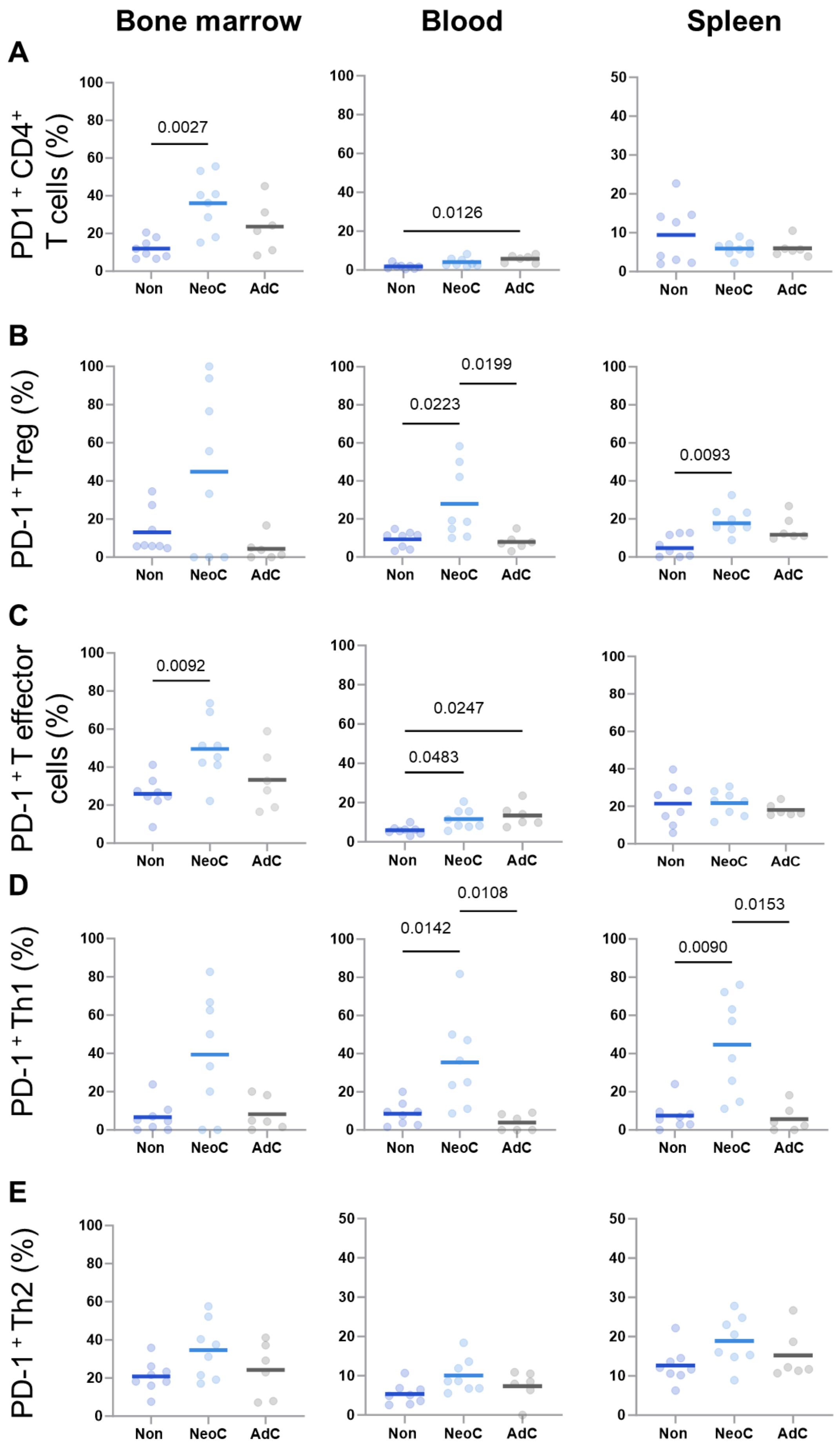
Bone marrow
Blood
Spleen
A
PD1 + CD4+ T cells (%)
0.0027
0.0126
B
PD-1 + Treg (%)
0.0199
0.0223
0.0093
C
PD-1 + T effector cells (%)
0.0092
0.0247
0.0483
D
PD-1 + Th1 (%)
0.0108
0.0142
0.0153
0.0090
E
PD-1 + Th2 (%)
Non
NeoC
AdC

**Figure S8. PD1$^+$ T cell subgroups in bone marrow, blood and spleen.** PD1$^+$ **A)** CD4$^+$ T cells **B)** T regulatory (Treg), **C)** T effector cells, **D)** T helper 1 (Th1) and **E)** T helper 2 (Th2) cells in bone marrow, blood and spleen between non-colonized (Non: n=8), neonatal-colonized (NeoC: n=8) and adult-colonized (AdC: n=6) groups from individual mice shown as dots with mean as line. Statistical analyses performed using one-way ANOVA with Tukey's multiple comparisons test or non-parametric with Kruskal-Wallis test. Only significant p-values shown.

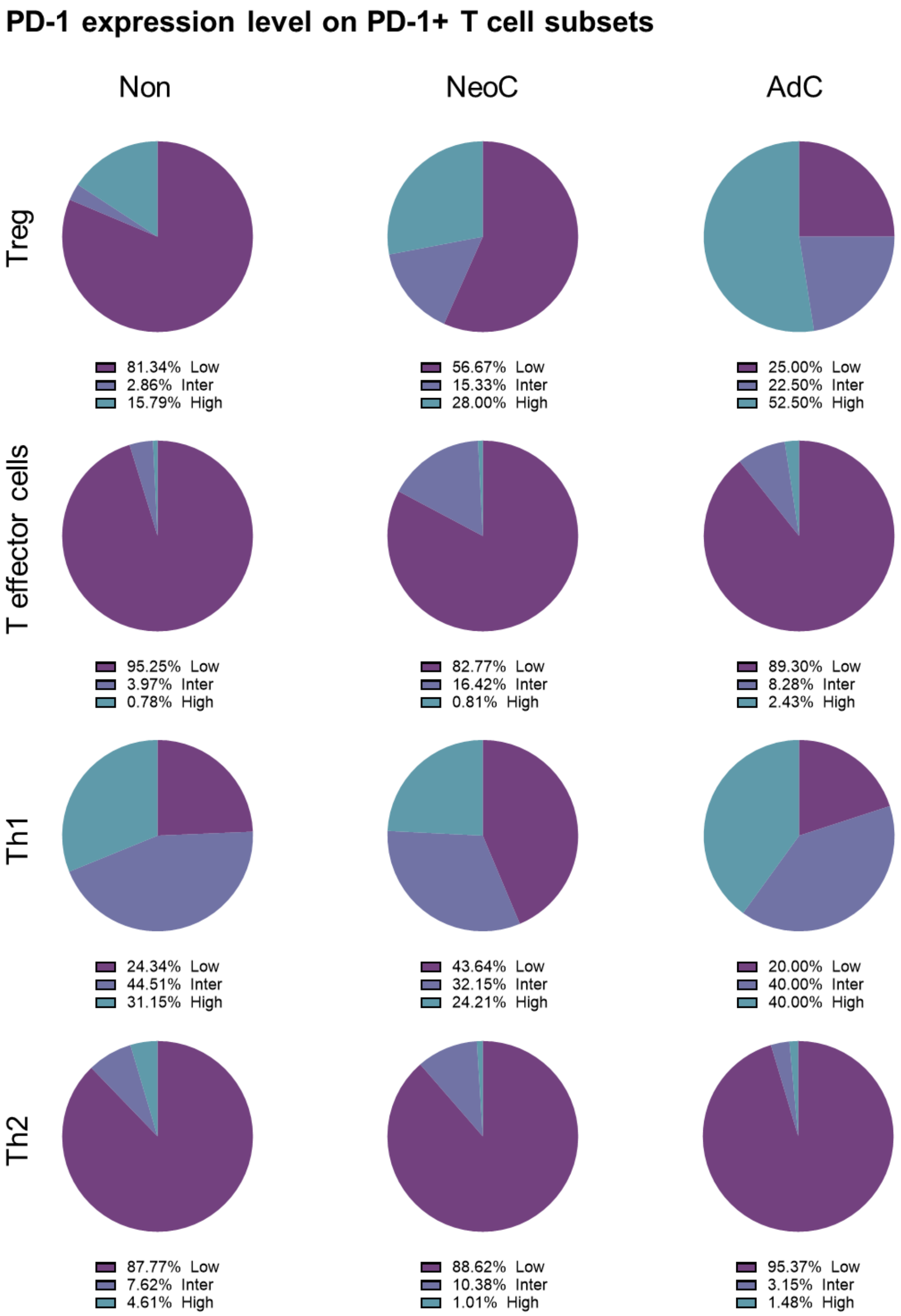


**Figure S9. PD-1 expression levels on PD-1$^+$ T cell subsets in the bone marrow.** PD-1 expression is shown as average of all mice from within either non- (Non), neonatal- (NeoC) or adult- (AdC) colonized groups on **A)** T regulatory (Treg), **B)** T effector **C)** T helper 1 (Th1) and **D)** T helper 2 (Th2) cells. The percentage of low, intermediate (Inter) or high PD-1 expression of the total PD-1$^+$ T cells within a subtype is indicated below each pie chart. .

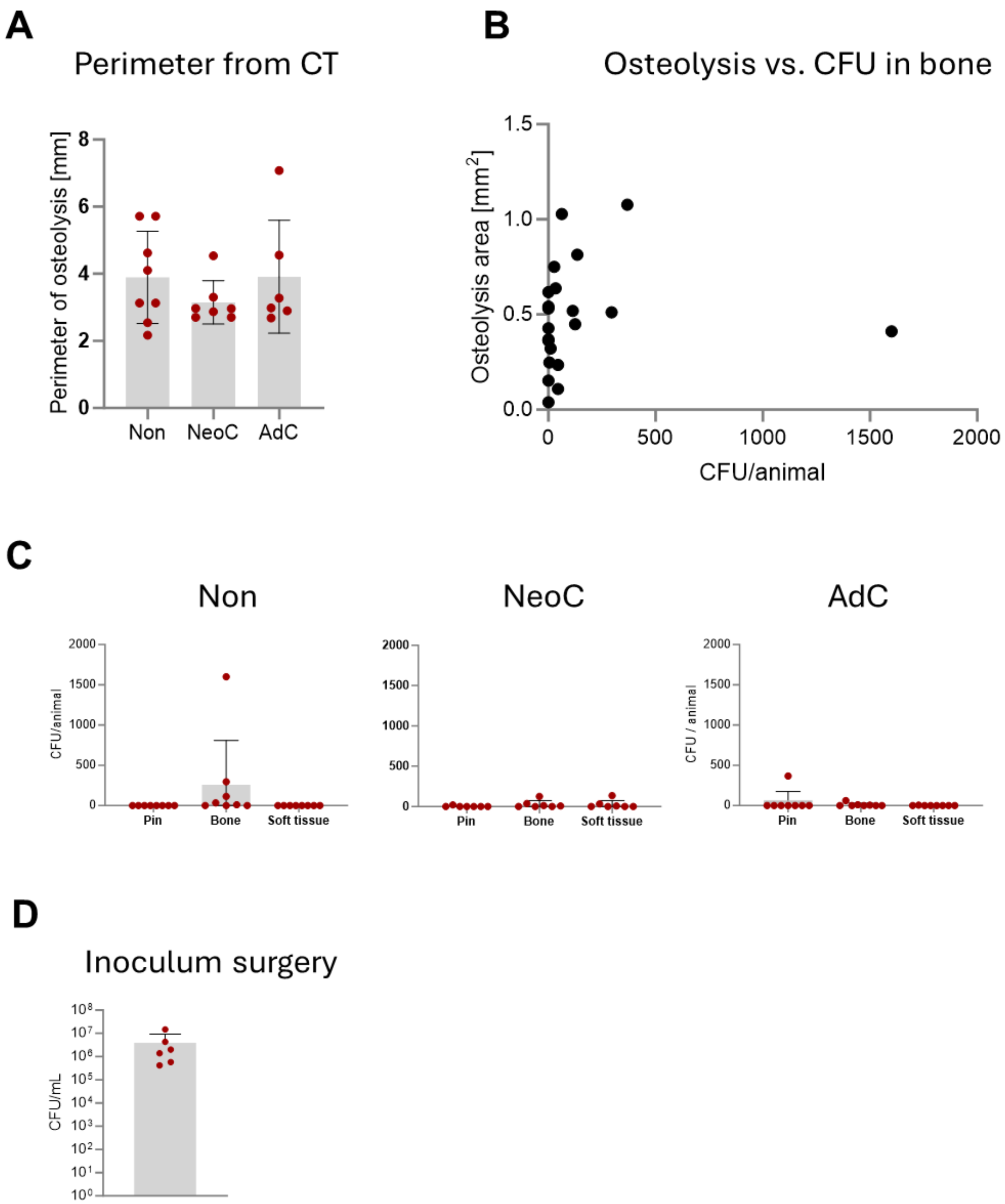


**Figure S10. Bacterial quantification and osteolysis compared between colonization groups. A)** Micro CT was performed at time of euthanasia. Osteolysis was identified and the perimeter of osteolysis area was measured. Each dot represents one mouse with the standard deviation in black and the mean as grey bar for each group (Non, n=8; NeoC, n=8; AdC, n=6). **B)** For each animal, the area of osteolysis was plotted against the CFU count of all tissues combined. There is no clear correlation. **C)** Following euthanasia, the implant (pin), the tibia (bone) and the surrounding soft tissue were harvested from each mouse and prepared for CFU/animal quantification. The results are divided per tissue and non- (Non; n=8), neonatal- (NeoC; n=8) and adult-colonization (AdC; n=8) groups and shown as dots with black standard deviation and grey mean. **D)** Bacterial colony forming units (CFU) per mL were quantified for one pin each per surgery day, shown as dots with black standard deviation and grey mean.

### Supplementary Material and Methods

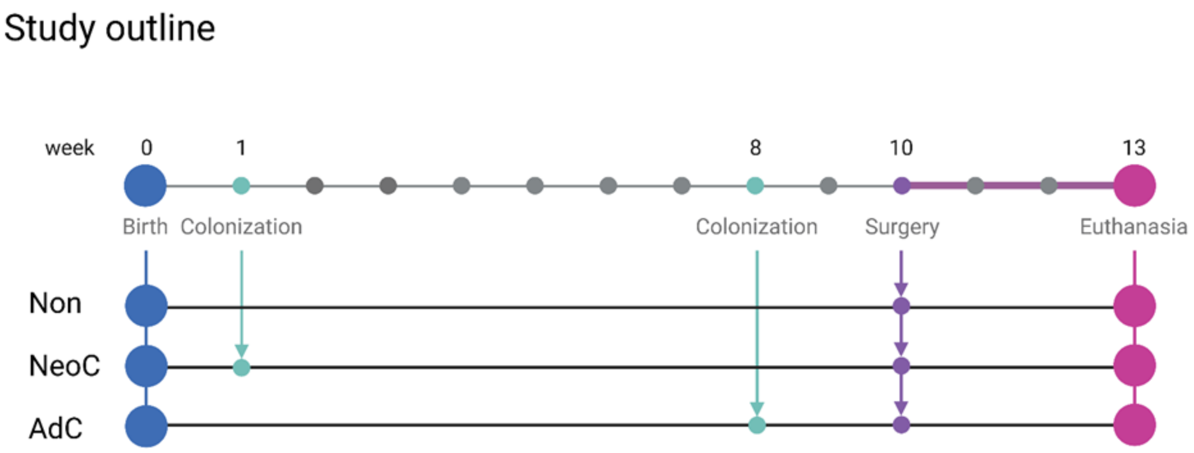


**Figure S11. Study outline and timeline:** Mice were either not colonized (Non), neonatal-colonized (NeoC) at 1 week of age or adult-colonized (AdC) at 8 weeks of age. All three groups received surgery, with a titanium pin inoculated with *Staphylococcus epidermidis* into their right tibia from anterior to posterior at week 10. Mice were euthanized 3 weeks later, at age of 13 weeks. Mice from each group were used for flow cytometry analysis of immune cells from the tibia bone marrow, the blood or the spleen, colony forming unit (CFU) determination to see the bacterial burden from homogenized tibia, the surrounding soft tissue and the pin, or for histology. Created in https://BioRender.com.

<u>Inoculum preparation</u>

*S. epidermidis* was adjusted to $OD_{600}$ of 5. Titanium pins (0.2 x 0.5 mm cross-section, 4 mm length, L-shape with 1 mm bend) were incubated in the bacterial suspension for 20 min, air-dried for 5 min in a sterile hood, and used within 4 h. One pin per batch was placed in 1 mL PBS, sonicated for 3 min, serially diluted, and plated on Tryptic soy agar (TSA, Oxiod, Basel, Switzerland) to determine colony forming units (CFU). Target bacterial load was $10^6$-$10^7$ CFU per pin. An inoculated pin was surgically implanted into the right proximal tibia of each mouse.

<u>Anesthesia, surgery and postoperative care</u>

Tramadol (100 mg/L) was added to drinking water the day before surgery and for 3 days postoperatively. Cages were changed 1–2 days before surgery and placed in a 27° C warming cabinet on the morning of surgery. Anesthesia was induced and maintained using sevoflurane (Baxter, Unterschleissheim, Germany) in oxygen, and buprenorphine (0.1 mg/kg sc, Bupaq, Streuli Tiergesundheit, Uznach, Switzerland) was given as intraoperative analgesia. Warmed Ringer's lactate (1 mL sc) was administered pre-surgery, and temperature was maintained via heating mat.
The right hindleg was aseptically prepared, and the implant site (2-3 mm below the tibial plateau) was identified using the proximal patella as an anatomical landmark. A hole was pre-drilled percutaneously through the proximal tibia from the medial to lateral cortex using a 25-gauge needle. The *S. epidermidis*-

inoculated pin was then inserted through the pre-drilled hole, leaving the bent end subcutaneous. If necessary, the puncture was sutured with simple interrupted stitches (Vicryl 5-0, C-3). Post-surgery, mice were returned to home cages with prior cage mates and kept in the warming cabinet until evening. Recovery food (DietGel® Recovery, ClearH2O, Westbrook, USA) and food soaked in tramadol water were provided on the cage floor. Animals were monitored twice daily for 5 days, daily for 2 days and twice weekly thereafter using a predefined score sheet assessing behavior, lameness, wound healing, Mouse Grimace Scale, respiration, outer appearance (fur), feces and weight. All mice were euthanized 3 weeks after surgery (13 weeks of age) by intracardial pentobarbital injection (Eskonarkon, Streuli Tiergesundheit, Uznach, Switzerland) under deep sevoflurane anesthesia. Six mice were excluded due to misplaced pins, high blood loss, or a postoperative fracture.

Flow cytometry staining

Table S3 lists antibodies used for surface staining of innate immune cells and B cells. The amount was titrated for optimal staining in our laboratory.

*Table S3. Panel innate immune cells and B cells*

| **Name** | **Amount/ 100uL** | **Clone** | **Lot number** | **Color** |
|---|---|---|---|---|
| Ly6G | 1 | 1A8 | B314454 | BV 650 |
| NK1.1 | 1 | PK136 | 2749831 | BV 605 |
| CD45 | 3 | 30-F11 | B322199 | BV510 |
| CD20 | 3 | QCH6A7 | 2749828 | BV421 |
| Via | 0.5 | 7AAD | - | FITC |
| CD3 | 2 | 145-2C11 | B307927 | PerCyP-Cy5.5 |
| CD80 | 2 | 16-10A1 | B386839 | PE |
| F4/80 | 3 | BM8 | 2442140 | PE-TexasRed |
| CD11c | 3 | N418 | 2518313 | PE-Cy5 |
| CD11b | 3 | M1/70 | B279418 | APC |
| CD19 | 2 | 6D5 | B363133 | Alexa-Fluor700 |
| Ly-6C | 2 | HK1.4 | B375238 | APC Cy7 |

Table S4 lists antibodies used for surface staining as well as intracellular staining (marked in grey) for T cells. The amount was titrated to the lowest sufficient amount in our laboratory.

*Table S4. Panel T cells subsets*

| **Name** | **Amount/ 100uL** | **Clone** | **Lot number** | **colour** |
|---|---|---|---|---|
| CD3 | 2 | 17A2 | B282105 | BV 650 |
| PD-1 | 1 | RNMP1-30 | 2547856 | BV605 |
| CD44 | 1 | IM7 | 2728724 | BV421 |
| dump channel (788AD/<br>CD11b/<br>Ly6C) | 0.5<br>3<br>2 | 7AAD<br>HK1.3<br>M1/70 | -<br>B390809<br>B389533 | FITC |
| Peptid (*S. epidermidis* 2w) | 1 | - | - | PE |
| CD8a | 5 | 53-6.7 | B278296 | PE-TexasRed |
| CTL-A4 | 3 | UC10-4B9 | 2519401 | PE-Cy7 |
| Fas (CD95) | 2 | SA367H8 | B364325 | APC |
| CD4 | 2 | GK 1.5 | B313203 | Alexa-Fluor700 |
| ICOS | 3 | C398.4A | B345307 | APC-Cy7 |
| IL-17a | 3 | eBio17B7 | 2749832 | BV 786 |
| IFN-gamma | 2 | XMG1.2 | 2687451 | BV 711 |
| IL-4 | 2 | 11B11 | 2666835 | PerCP |
| FOXP3 | 2 | FJK-16s | 2607676 | PE-Alexa Fluor700 |

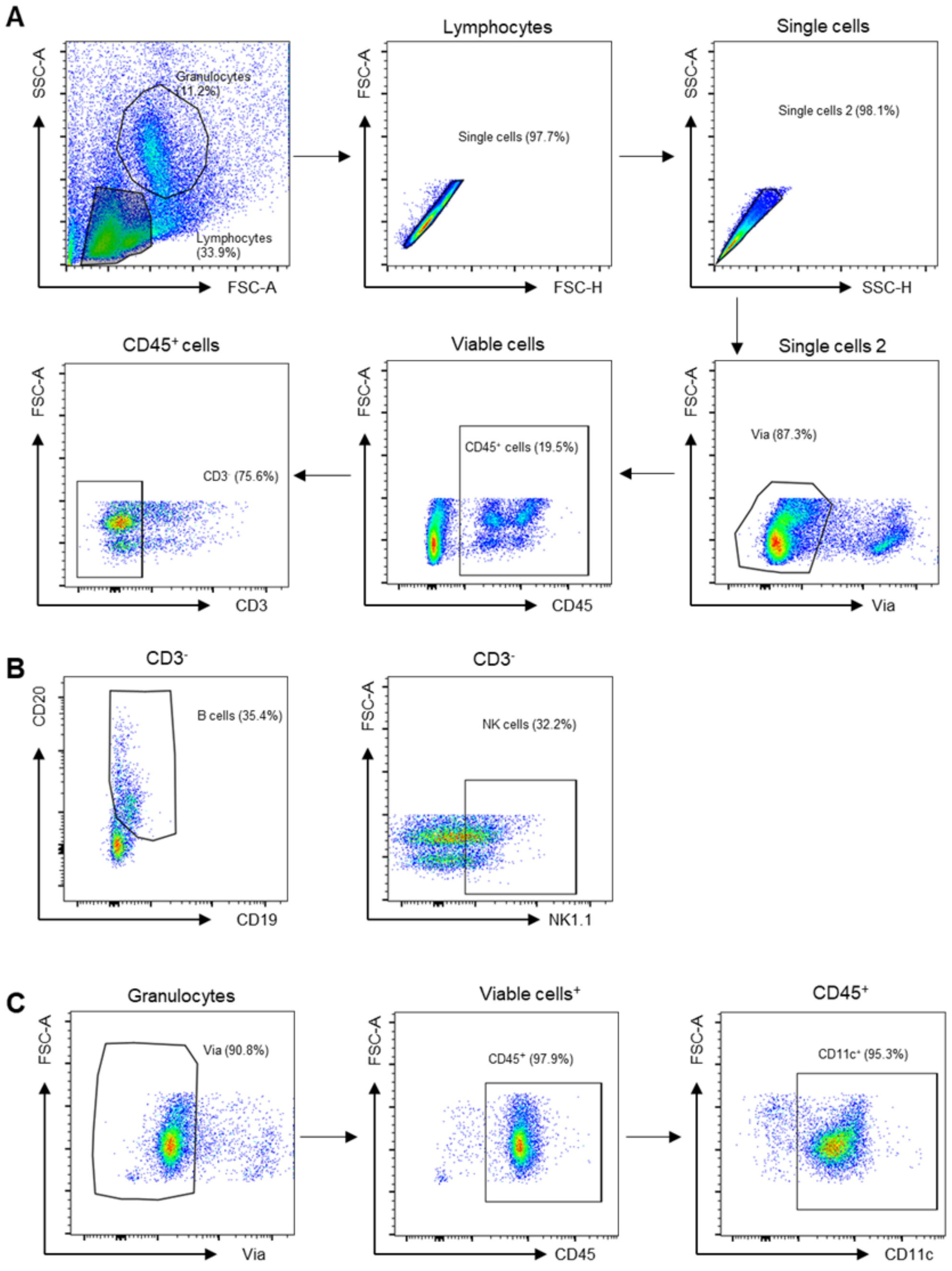


**Gating strategy innate and B cells**

**Figure S12. Flow cytometry gating strategy of innate immune cells and B cells A)** Flow cytometry gating strategy for the identification and classification of innate immune cell populations and B cells from bone marrow, blood and spleen. Titles represent the gates that were gated on. All representative scatter plots presented are obtained from bone marrow samples of non-colonized mice. The data were analyzed using FlowJo software V10. First, Lymphocytes and granulocytes were gated (FSC-A against SSA-A). Next, doublet cells were excluded (FSC-A against FSC-H and SSC-A against SSC-H). Dead cells were excluded (Via against FSC-A), CD45$^+$ were gated (CD45 against FSC-A) and CD3$^+$ cells excluded (CD3 against FSC-A). **B)** On CD3$^+$ cells, B cells (CD19 against CD20) and Natural Killer cells (NK1.1 against FSC-A) were gated. **C)** On granulocytes, dead cells were excluded (Via against FSC-A), CD45$^+$ were gated (CD45 against FSC-A) and CD11c$^+$ cells gated (CD11c against FSC-A).

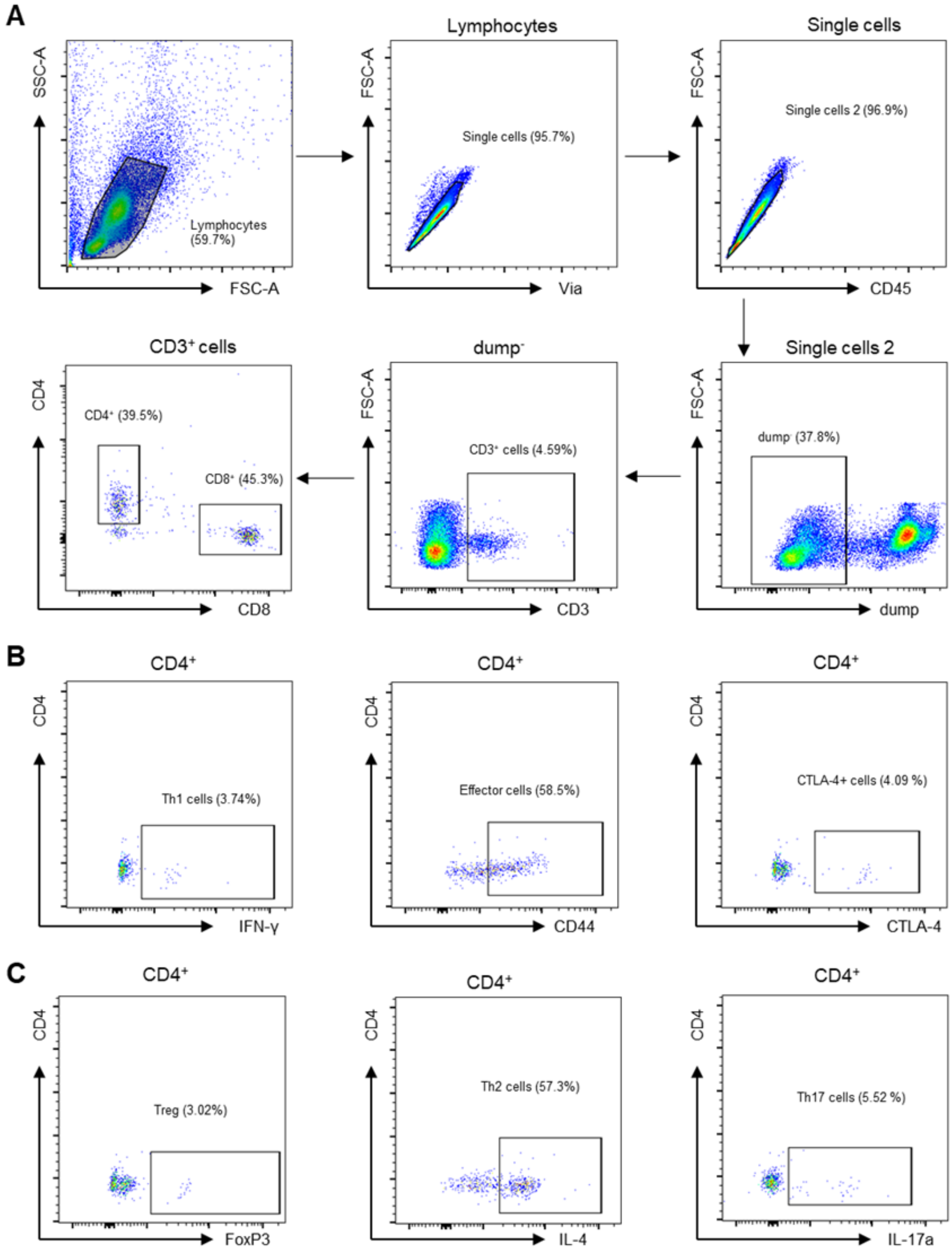


**Gating strategy T cells**

**Figure S13. Flow cytometry gating strategy of T cell subsets. A)** Flow cytometry gating strategy for the identification and classification of T cell subsets from bone marrow, blood and spleen. Titles represent the gates that were gated on. All representative scatter plots presented are obtained from bone marrow samples of non-colonized mice. The files were analyzed using FlowJo software V10. First, Lymphocytes were gated (FSC-A against SSA-A). The next step was to exclude doublet cells (FSC-A against FSC-H and SSC-A against SSC-H). Dead cells as well as Ly6c$^+$, CD11c$^+$, CD19$^+$ cells were excluded in the dump channel. CD3$^+$ were gated (CD3 against FSC-A). CD8$^+$ as well as CD4$^+$ cells were gated (CD8 against CD4). **B)** On CD4$^+$ cells, Th1 (IFN-γ against CD4), Th2 (IL-2 against CD4) and Th17 cells (IL-17a against CD4). **C)** Treg (FoxP3 against CD4) as well as effector cells (CD44 against CD4) were gated on CD4 T cells.

Antibodies used for Immunofluorescence

Primary antibodies included goat anti-CD3-epsilon (clone M-20, sc-1127, RRID: AB_631128, Santa Cruz Biotechnology) and rabbit anti-*S. aureus* (PA1-7246, RRID:AB_561546, Thermo Fisher Scientific) used at a 1:100 dilution; rabbit-anti Tbet/Tbx21 (MBS248480, MyBiosource), rabbit anti-PD1 (Clone EPR20665, ab214421, RRID:AB_2941806, Abcam), biotin rat anti-Ly6G (clone 1A8, 127604, RRID: AB_1186108, Biolegend), rat anti-mouse Foxp3 (clone FJK-16S, 14-5773-82, RRID:AB_467576, Thermo Fisher Sceintific) and rat anti-mouse Ki67 (HS-398 117, HistoSure) used at a 1:50 dilution. Secondary antibodies were Alexa Fluor 568-conjugated donkey anti-goat IgG (A-11057, RRID: AB_2534104, Thermo Fisher Scientific) at 1:200 for CD3-epsilon, Alexa Fluor 488-conjugated donkey anti-rabbit IgG (711-546-152, RRID: AB_2340619, Jackson ImmunoResearch Laboratories) at 1:200 for S. aureus, Tbet and PD1; and Alexa Fluor 647 donkey anti-rat IgG (712-606-153, RRID: AB_2340696, Jackson ImmunoResearch Laboratories) at 1:200 for Foxp3.

Sections were incubated at 60 °C overnight for deparaffinization, transferred to xylene, and rehydrated through graded ethanol (100%, 96%, 70%) to water. Antigen retrieval was performed in Antigen Unmasking Solution (S1699, Agilent) by boiling for 2 h. Nonspecific binding was blocked with 5% donkey serum in Phosphate-Buffered Saline (PBS, MT21040CV, Thermo Fisher Scientific) containing 0.5% Triton X-100 for 40 min in a humidified chamber.
